# SAMHD1 enhances HIV-1-induced apoptosis in monocytic cells via the mitochondrial pathway

**DOI:** 10.1101/2025.01.08.632057

**Authors:** Hua Yang, Pak-Hin Hinson Cheung, Li Wu

## Abstract

Sterile alpha motif (SAM) and histidine-aspartate (HD) domain-containing protein 1 (SAMHD1) inhibits HIV-1 replication in non-dividing cells by reducing the intracellular dNTP pool. SAMHD1 enhances spontaneous apoptosis in cells, but its effects on HIV-1-induced apoptosis and the underlying mechanisms remain unknown. Here we uncover a new mechanism by which SAMHD1 enhances HIV-1-induced apoptosis in monocytic cells through the mitochondrial pathway. We found that endogenous SAMHD1 enhances apoptosis levels induced by HIV-1 infection in dividing THP-1 cells. Mechanistically, SAMHD1 expression decreases the mitochondrial membrane potential and promotes cytochrome c release induced by HIV-1 infection in THP-1 cells, thereby enhancing mitochondrial apoptotic pathway. SAMHD1-enhanced apoptosis is associated with increased expression of the pro-apoptotic protein BCL-2-interacting killer (BIK) in cells. We further demonstrated that BIK contributes to SAMHD1-enhanced apoptosis during HIV-1 infection. Overall, our results reveal an unappreciated regulatory mechanism of SAMHD1 in enhancing HIV-1-induced apoptosis via the mitochondrial pathway in monocytic cells.

## INTRODUCTION

As a dNTP triphosphohydrolase that reduces intracellular dNTP levels,^1,2^ SAMHD1 restricts the reverse transcription of retroviruses, including human immunodeficiency virus type 1 (HIV-1)^1,3,4^ and certain simian immunodeficiency viruses (SIV).^5^ SAMHD1 restricts HIV-1 infection in non-diving immune cells, such as resting CD4+ T cells, macrophages, and dendritic cells. ^3,4,6^ Through different mechanisms, SAMHD1 also restricts the replication of DNA viruses, including human papilloma virus,^7^ hepatitis B virus,^8,9^ and human cytomegalovirus,^10–12^ as well as RNA viruses, including enteroviruses^13,14^ and influenza A virus^15,16^. We have reported that SAMHD1 suppresses NF-κB activation and type I interferon (IFN-I) activation induced by virus infections and inflammation through interaction with the key proteins in the NF-κB and IFN-I pathways.^17–19^ Multifaceted SAMHD1 also regulates a variety of cellular processes,^20–22^ including DNA replication fork progression,^23^ cell proliferation^7^ and apoptosis^24,25^. However, the mechanisms by which SAMHD1 regulates these processes remain unclear.^20^

Apoptosis is a major form of programmed cell death and is divided into death receptor apoptotic pathway and mitochondrial apoptotic pathway.^26,27^ In the death receptor apoptotic pathway, certain ligands, such as Fas ligand (Fas-L), bind to death receptors, leading to recruitment and activation of cysteine-aspartyl proteases (caspase) 8. Activated caspase 8 cleaves and activates downstream of caspases 3 and 7 (3/7) as executioners, which then cleave poly (ADP-ribose) polymerase (PARP), ultimately triggering apoptosis. In the mitochondrial apoptotic pathway, death receptor signals or mitochondrial apoptotic stimuli, including DNA damage and mitotic arrest, trigger the mitochondrial outer membrane permeabilization (MOMP). This results in a decrease in the mitochondrial membrane potential (Δψm) and the release of cytochrome c (Cyto c) from the mitochondria into the cytosol. In the cytosol, Cyto c forms apoptosome with apoptotic peptidase activating factor 1, which activates the caspase cascade. Specifically, initiator caspase 9 is cleaved and activated, leading to the activation of caspases 3/7 to cleave a large set of substrates that ultimately lead to apoptotic cell death.

Specific substrates of caspases 3/7 have been used as markers to indicate apoptosis, such as cleavage of PARP (or PARP1). The mitochondrial apoptotic pathway is tightly regulated by the B cell lymphoma 2 (BCL-2) protein family, which contains conserved BCL-2 homology (BH) domains.^28^ The BCL-2 protein family is divided into three subsets based on apoptotic function: anti-apoptotic BCL-2 proteins, including BCL-2, and B cell lymphoma extra-large (BCL-X_L_), apoptotic effectors, including BCL-2 associated X (BAX) and BCL2 antagonist/killer (BAK), and pro-apoptotic BH3-only proteins, including BH3-interacting domain death agonist (BID), BCL-2 interacting mediator of cell death (BIM), and BCL2 interacting killer (BIK). The apoptotic effectors oligomerize and form pores in the outer mitochondrial membrane to induce MOMP. Anti-apoptotic BCL-2 proteins interact with apoptotic effectors to neutralize the change of MOMP, while BH3-only proteins interact with anti-apoptotic proteins BCL2, disrupting their inhibition of BAX/BAK, thereby promoting apoptosis.^28^

HIV-1 infected individuals without effective antiretroviral therapy experience a progressive decrease in lymphocyte count, particularly CD4+ T cells.^29^ HIV-1 induces cell death primarily through apoptosis,^30^ pyroptosis,^31,32^ necrosis^33^, and necroptosis.^33,34^ HIV-1 infection induces apoptosis through multiple viral proteins, including envelope glycoprotein (gp120), transcription activator (Tat), negative regulatory factor (Nef), and viral protein regulatory (Vpr).^35,36^ HIV-1 Vpr blocks the cells cycle at the G2 phase, which promotes apoptosis, and disrupts MOMP to facilitate the release of Cyto c.^37,38^

SAMHD1 restricts human T cell leukemia virus type 1 (HTLV-1) infection of primary monocytes by accumulating viral reverse transcription intermediates, which activate the DNA sensing pathway and lead to apoptosis.^39^ We have demonstrated that exogenous SAMHD1 expression enhances Fas-L induced apoptosis through death receptor apoptotic pathway in T-cell lymphoma-derived HuT78 cells.^40^ We have also reported that endogenous SAMHD1 enhances spontaneous apoptosis in THP-1 cells.^24^ However, the function and mechanisms of SAMHD1 in regulating HIV-1-induced apoptosis remains unclear.

In this study, we found that endogenous SAMHD1 enhances apoptosis induced by HIV-1 infection in THP-1 cells through the mitochondrial pathway. We demonstrated that SAMHD1 decreased Δψm and increased the release of Cyto c from the mitochondria into the cytosol in HIV-1-infected cells. Furthermore, SAMHD1-enhanced apoptosis was associated with increased BIK expression and BIK contributed to SAMHD1-enhanced apoptosis during HIV-1 infection. Our findings reveal a regulatory mechanism by which SAMHD1 enhances apoptosis induced by HIV-1 infection, suggesting a previously unappreciated function of SAMHD1 in modulating cellular responses to viral infection.

## RESULTS

### Endogenous SAMHD1 enhances apoptosis induced by HIV-1 infection of THP-1 cells

Our previous study demonstrated that SAMHD1 inhibited single-cycle HIV-1 infection in dividing THP-1 cells at 1 day post infection (dpi), but not at 2 dpi.^24^ Similar HIV-1 infection levels at 2 dpi or later in dividing THP-1 cells with or without endogenous SAMHD1 provide an appropriate model to investigate the function of SAMHD1 in HIV-1-induced apoptosis in monocytic cells. ^24^ To examine the effects of SAMHD1 on HIV-1 infection in monocytic THP-1 cell line, THP-1 control (Ctrl) and SAMHD1 knock-out (KO) cell lines were infected with a single-cycle luciferase reporter HIV-1 pseudotyped with VSV-G (HIV-1-Luc/VSV-G), which lacks viral envelope protein and Nef. ^24^ The HIV-1 reverse transcriptase inhibitor nevirapine (NVP) was used to treat infected cells to block HIV-1 replication. Luciferase activity, which indicates HIV-1 replication, was measured at 1 to 4 dpi. At 1 dpi, we detected higher luciferase activity in infected SAMHD1 KO cells compared to Ctrl cells, supporting that SAMHD1 inhibits HIV-1 replication at early time in THP-1 monocytic cells (Figure 1A). At 2 and 3 dpi, however, luciferase activity became comparable between infected Ctrl cells and SAMHD1 KO cells. Conversely, at 4 dpi, higher luciferase activity was detected in infected Ctrl cells compared to SAMHD1 KO cells, consistent with our previous results that SAMHD1 did not inhibit HIV-1 infection at later time points in monocytic cells.^24^ As expected, NVP treatment completely inhibited HIV-1 replication from 1 to 4 dpi (Figure 1A).

**Figure 1.**
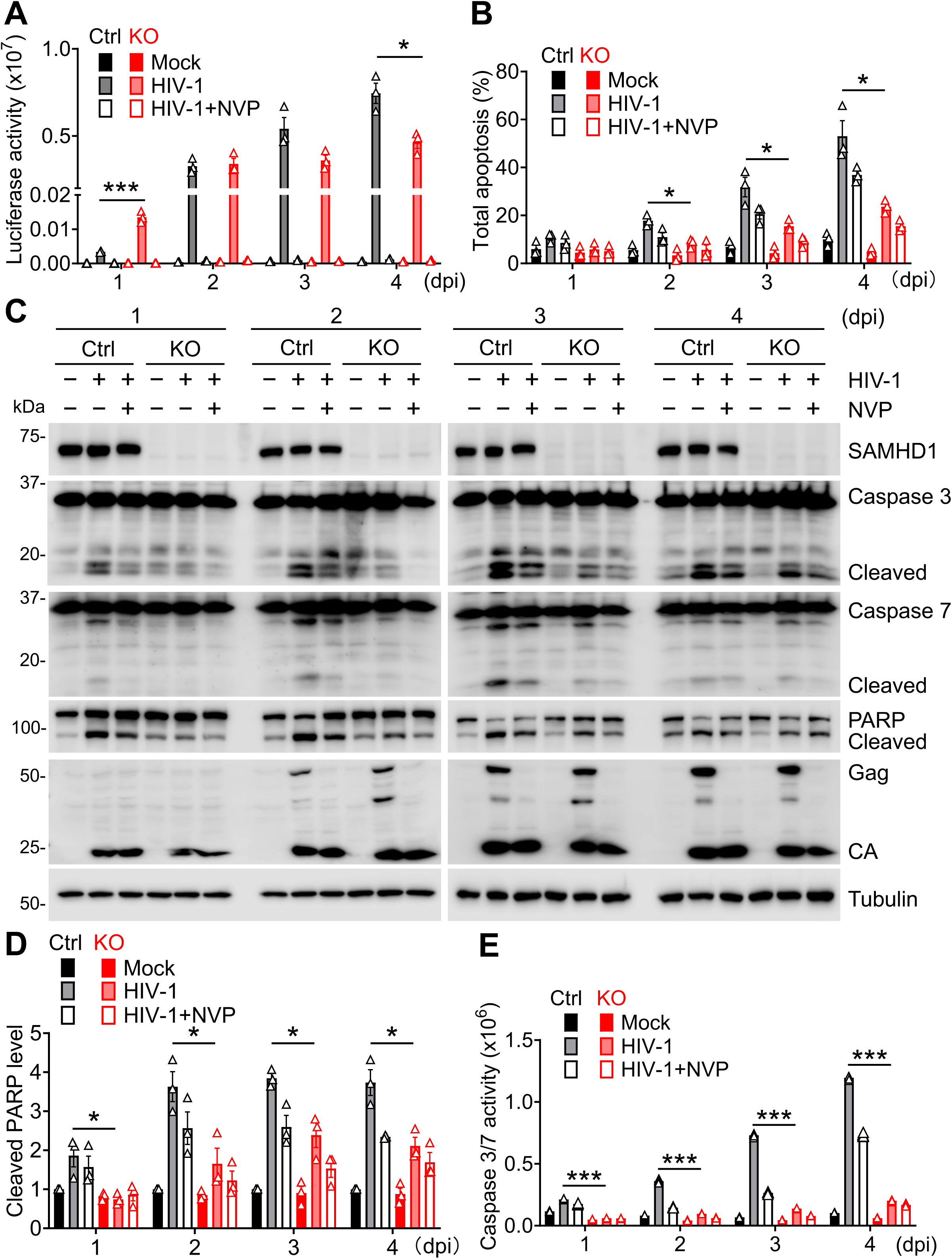
Endogenous SAMHD1 enhances apoptosis induced by HIV-1 infection of THP-1 cells. **(A-E)** THP-1 control (Ctrl) and stable SAMHD1 knock-out (KO) cell lines were infected with single-cycle, luciferase reporter HIV-1-Luc/VSV-G [multiplicity of infection=2 (MOI=2)] or mock infected. The HIV-1 reverse transcriptase inhibitor nevirapine (NVP) was used to treat infected cells to block HIV-1 infection. Cells were harvested at 1-4 days post infection (dpi) for analyses. **(A)** HIV-1 infection levels were measured by a luciferase assay and normalized for cellular protein concentration. **(B)** Apoptosis levels were measured by Annexin V and 7-AAD staining and flow cytometry. The percentages of total apoptosis include early apoptosis (positive for Annexin V but negative for 7-AAD) and late apoptosis (positive for both Annexin V and 7-AAD). Original flow cytometry results are shown in Fig. S1A. **(C)** Detection of SAMHD1, caspases 3/7, poly (ADP-ribose) polymerase (PARP), HIV-1 Gag and capsids (CA), and tubulin by Western blot. Tubulin was used as a loading control. The cleaved caspases 3/7 and PARP proteins are indicated. **(D)** The cleaved PARP levels were quantified by densitometry analysis and normalized to tubulin. The level of cleaved PARP of mock-infected THP-1 Ctrl cells was set to 1. **(E)** Caspase 3/7 activities were measured by the Caspase-Glo 3/7 assay. **(A-B and D-E)** Data are presented as means ± SEM. The results in A, B, and D represent three independent experiments and the results in E represent six independent experiments. The unpaired t-test was used for statistical significance compared with THP-1 Ctrl cells with HIV-1 infection. *p < 0.05; ***p < 0.001.

Our previous study showed that SAMHD1 enhances spontaneous apoptosis in THP-1 cells.^24^ To analyze the effects of SAMHD1 on HIV-1-induced apoptosis in the monocytic cells, THP-1 Ctrl and SAMHD1 KO cell lines were infected with HIV-1-Luc/VSV-G for 1 to 4 days. Annexin-V and 7-AAD staining was used to detect early apoptosis (positive for annexin-V but negative for 7-AAD) and late apoptosis (positive for both annexin-V and 7-AAD). Flow cytometry analysis revealed that the percentages of total apoptosis were higher in infected THP-1 Ctrl cells compared to infected SAMHD1 KO cells at 2 to 4 dpi, suggesting that SAMHD1 enhances HIV-1-induced apoptosis in monocytic cells (Figures 1B and S1A). NVP treatment partially inhibited HIV-1-induced apoptosis but did not affect spontaneous apoptosis (Figures 1B, S1A, and S2).

To investigate the mechanisms by which SAMHD1 enhances apoptosis in cells, cleaved caspases 3/7 (effector caspases) and cleaved PARP (alternatively PARP-1, an apoptosis hallmark) were detected by Western blot. THP-1 Ctrl cells showed enhanced cleavage of caspases 3/7 and PARP upon HIV-1 infection compared to SAMHD1 KO cells at 1 to 4 dpi (Figure 1C). The treatment of NVP partially inhibited HIV-1-induced cleavage of caspases 3/7 and PARP in both THP-1 Ctrl and SAMHD1 KO cells. Quantification of the cleavage of PARP revealed that HIV-1-induced PARP cleavage was stronger in THP-1 Ctrl cells than that in SAMHD1 KO cells at 1 to 4 dpi (Figure 1D). Furthermore, HIV-1-Luc/VSV-G-infected THP-1 Ctrl cells showed higher caspase 3/7 activity than SAMHD1 KO cells at 1 to 4 dpi (Figure 1E). NVP treatment partially inhibited HIV-1-induced caspase 3/7 activity. Thus, endogenous SAMHD1 enhances apoptosis induced by single-cycle HIV-1 infection in THP-1 cells.

### SAMHD1 reconstitution in THP-1 SAMHD1 KO cells enhances apoptosis induced by HIV-1 infection

To confirm the function of SAMHD1 in enhancing apoptosis, we reconstituted SAMHD1 expression in THP-1 SAMHD1 KO cells and assessed HIV-1-induced apoptosis. THP-1 Ctrl, SAMHD1 KO, Lvx vector control, and SAMHD1 knock-in (KI) cell lines were infected with HIV-1-Luc/VSV-G or mock infected for 2 and 3 dpi. THP-1 Lvx vector control cells and SAMHD1 KI cells were generated from the parental SAMHD1 KO cells^24^. HIV-1-Luc/VSV-G infection efficiency was similar across the four cell lines at 2 dpi and 3 dpi (Figure 2A). The percentages of total apoptosis in THP-1 Ctrl and SAMHD1 KI cells, which expressed similar levels of SAMHD1 protein, were higher than that observed in SAMHD1 KO and Lvx cells upon HIV-1-Luc/VSV-G infection (Figures 2B, 2C, and S1B). THP-1 Ctrl and SAMHD1 KI cells exhibited enhanced cleavage of caspases 3/7 and PARP compared to SAMHD1 KO and Lvx cells upon HIV-1-Luc/VSV-G infection (Figure 2C). Quantification of three independent experiments showed that the cleavage of PARP induced by HIV-1 infection was stronger in THP-1 Ctrl and SAMHD1 KI cells than SAMHD1 KO and Lvx cells at 2 and 3 dpi (Figure 2D). Consistently, THP-1 Ctrl and SAMHD1 KI cells had higher HIV-1-induced caspase 3/7 activity compared to that observed in SAMHD1 KO and Lvx cells (Figure 2E). Thus, SAMHD1 reconstitution in THP-1 SAMHD1 KO cells enhances apoptosis induced by single-cycle HIV-1 infection.

**Figure 2.**
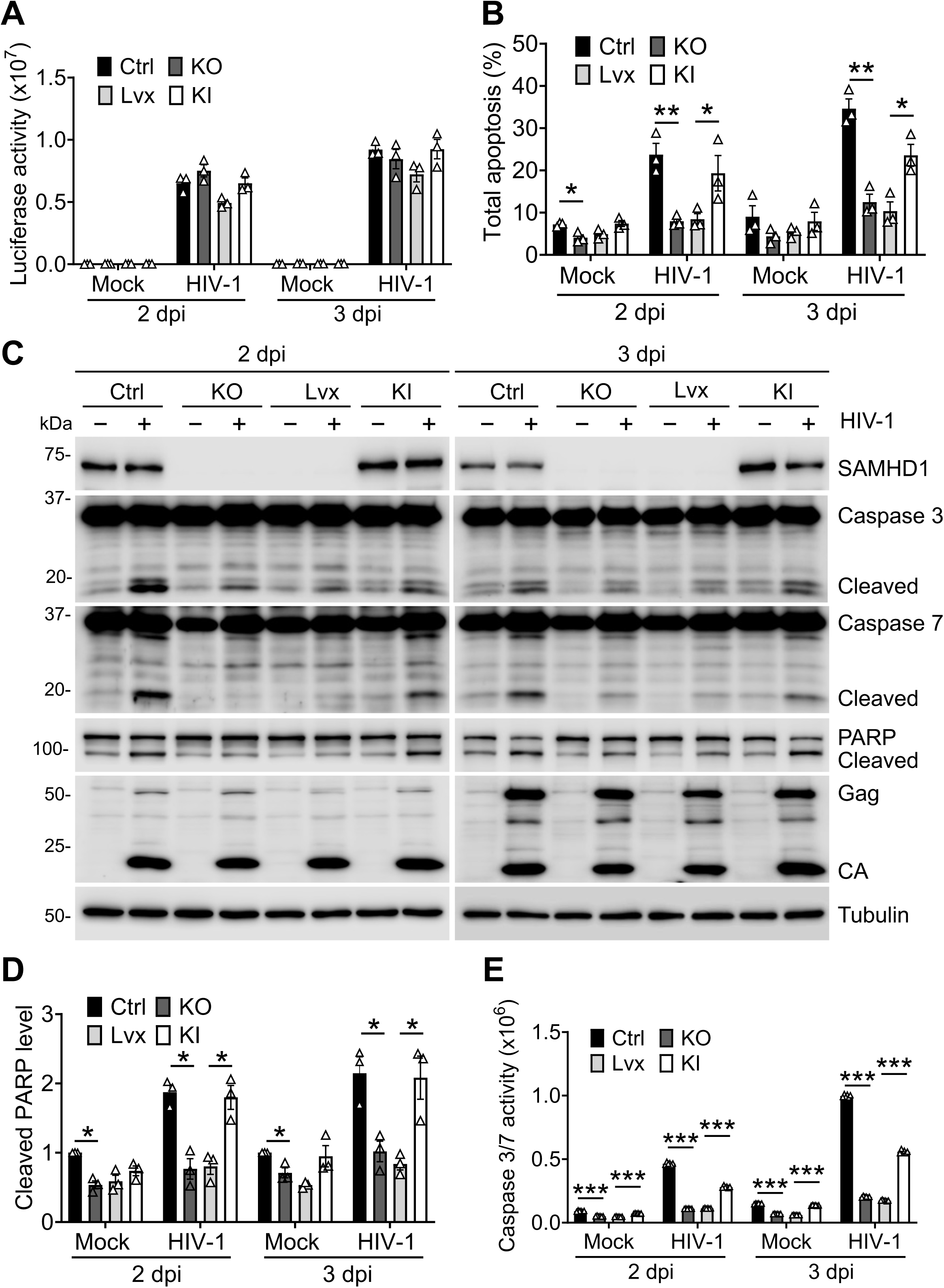
SAMHD1 reconstitution in THP-1 SAMHD1 KO cells enhances apoptosis induced by HIV-1 infection. **(A-E)** THP-1 Ctrl, SAMHD1 KO, Lvx vector control, and SAMHD1 knock-in (KI) cell lines were infected with HIV-1-Luc/VSV-G (MOI=2) or mock-treated. Cells were collected at 2 and 3 dpi for analyses. The data in A, B, and D (means ± SEM) represent three independent experiments and data in E represent six independent experiments. **(A)** The HIV-1 infection levels were measured by the luciferase assay and normalized for cellular protein concentration. **(B)** Total apoptosis of cells was measured by flow cytometry as indicated in Fig. 1B. Original flow cytometry results are shown in Fig. S1B. **(C)** Detection of SAMHD1, caspase 3, caspase 7, PARP, HIV-1 Gag and CA, and tubulin by Western blot. The cleaved caspases 3/7 and PARP proteins are indicated. **(D)** The relative cleaved PARP levels were quantified by densitometry analysis and normalized to tubulin. The level of cleaved PARP of mock-infected THP-1 Ctrl cells was set to 1. **(E)** Caspase 3/7 activities were measured by Caspase-Glo 3/7 assay. **(A-B and D-E)** The unpaired t test was used for statistical significance compared with THP-1 Ctrl cells or THP-1 Lvx cells. *p < 0.05; **p < 0.01; ***p < 0.001.

### NVP or Z-DEVD-FMK treatment inhibits SAMHD1-enhanced apoptosis induced by replication-competent HIV-1 infection

To investigate the role of SAMHD1 in apoptosis induced by wild-type replication-competent HIV-1 that expresses all viral proteins, THP-1 Ctrl and SAMHD1 KO cell lines were mock infected or infected with HIV-1_NL4-3_ in the presence or absence of NVP at 2, 4, 6 and 8 dpi. HIV-1_NL4-3_ infected THP-1 Ctrl cells generally exhibited higher apoptotic response than the infected SAMHD1 KO cells at 4 and 6 dpi as shown by quantification of the cleaved PARP levels (Figure 3A). Compared to SAMHD1 KO cells, HIV-1_NL4-3_ infection in THP-1 Ctrl cells induced higher cleavage of caspases 3/7 and PARP at 4 and 6 dpi (Figures 3A and S3A). Expression of HIV-1_NL4-3_ Gag (p55), immature Gag (p41), and capsid (CA) were similar between HIV-1-infected THP-1 Ctrl cells and SAMHD1 KO cells (Figure S3A). NVP treatment inhibited HIV-1_NL4-3_ replication and HIV-1-induced apoptosis in both THP-1 Ctrl and SAMHD1 KO cells (Figures 3A and S3A). Importantly, NVP treatment blocked HIV-1_NL4-3_ induced PARP cleavage in THP-1 Ctrl cells at 4, and 6 dpi (Figure 3A). Therefore, SAMHD1 promotes apoptosis induced by replication-competent HIV-1 infection in dividing THP-1 cells.

**Figure 3.**
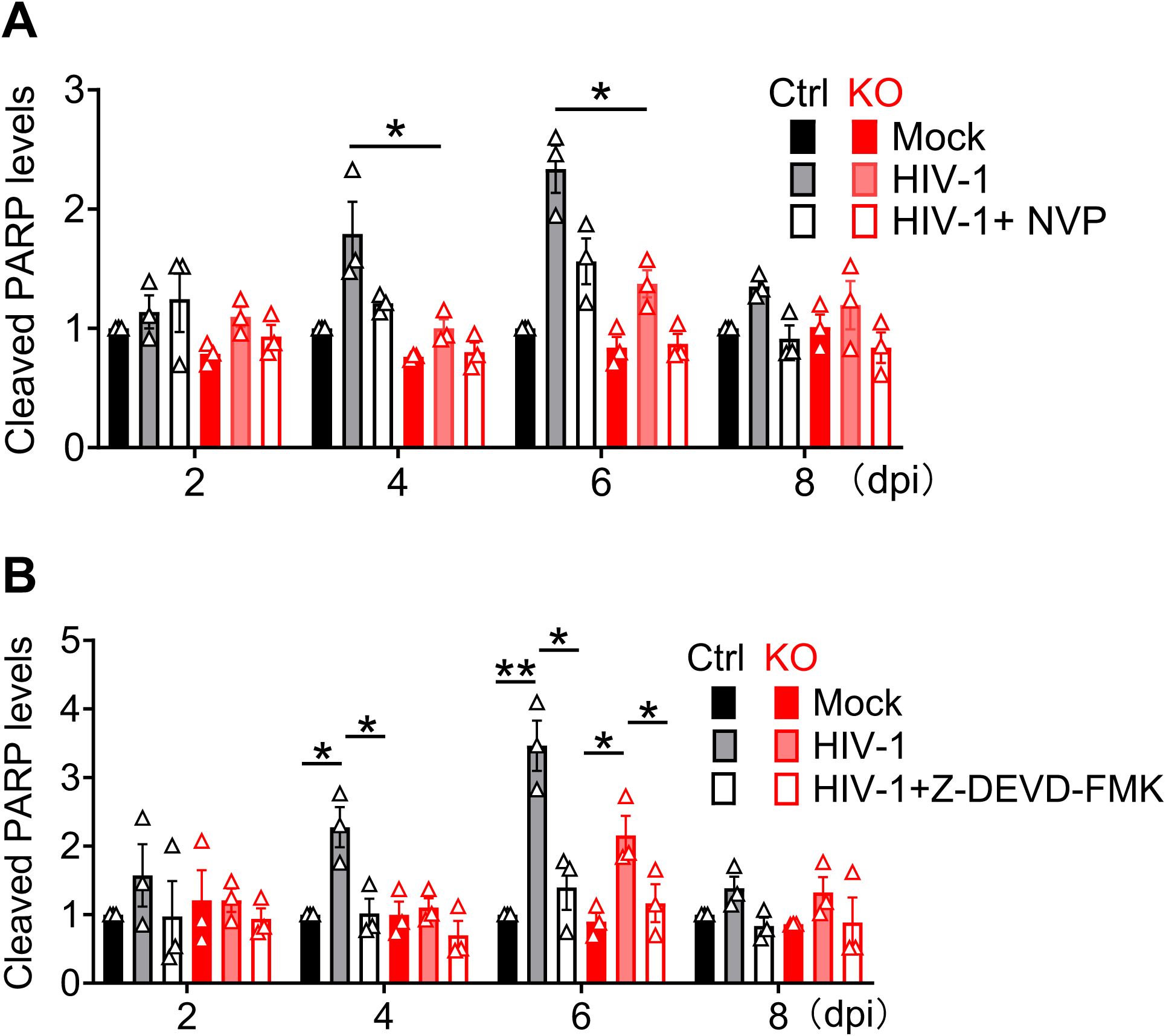
NVP or Z-DEVD-FMK treatment inhibits SAMHD1-enhanced apoptosis induced by replication-competent HIV-1 infection. **(A-B)** THP-1 Ctrl and SAMHD1 KO cell lines were infected with replication-competent HIV-1_NL4-3_ (MOI=2) or mock-treated, and harvested at 2, 4, 6, and 8 dpi for cleaved PARP analysis by Western blot. The relative cleaved PARP levels were quantified by densitometry analysis and represented three independent experiments. The level of cleaved PARP of mock-infected THP-1 Ctrl cells was set to 1. The unpaired t-test was used for statistical significance compared with THP-1 Ctrl cells and SAMHD1 KO cells with HIV-1_NL4-3_ infection. *p < 0.05; **p < 0.01. **(A)** NVP was used to block HIV-1 infection. Original Western blots are shown in Fig. S3A. **(B)** Z-DEVD-FMK was used to inhibit caspase 3/7 activity. Original Western blots are shown in Fig. S3B.

To examine whether SAMHD1-enhanced apoptosis by HIV-1 infection is dependent on caspase 3/7 activity, THP-1 Ctrl and SAMHD1 KO cell lines were infected or mock-infected with HIV-1_NL4-3_ and treated with Z-DEVD-FMK, which is a specific and irreversible caspase-3/7 inhibitor and widely used for inhibiting cell apoptotic death.^41,42^ Z-DEVD-FMK treatment did not affect the expression of HIV-1 Gag, immature Gag, and CA that were comparable between THP-1 Ctrl cells and SAMHD1 KO cells (Figure S3B), suggesting similar HIV-1_NL4-3_ replication efficiency in these cells. Z-DEVD-FMK treatment significantly mitigated HIV-1_NL4-3_ induced PARP cleavage in THP-1 Ctrl cells at 4 and 6 dpi (Figures 3B and S3B). HIV-1_NL4-3_ infection did not significantly induce PARP cleavage in SAMHD1 KO cells at 2, 4, and 8 dpi. Z-DEVD-FMK treatment also decreased HIV-1-induced cleavage of caspases 3/7 and PARP in THP-1 Ctrl at 4 and 6 dpi and in SAMHD1 KO cells at 6 dpi (Figures 3B and S3B). These data suggest that caspase 3/7 activities contribute to SAMHD1-enhanced apoptosis induced by HIV-1 replication in dividing THP-1 cells.

### SAMHD1 enhances apoptosis induced by HIV-1 infection through the mitochondrial pathway

To investigate whether SAMHD1 enhances apoptosis induced by HIV-1 infection through the mitochondrial apoptotic pathway, we measured Δψm of cells during HIV-1 infection. THP-1 Ctrl and SAMHD1 KO cell lines were mock-infected or infected with single-cycle HIV-1-Luc/VSV-G and treated with NVP. At 1, 2, 3 and 4 dpi, cells were stained by JC-1 to detect Δψm using flow cytometry. JC-1 is a fluorescent dye that serves as an indicator of Δψm during apoptosis. ^43^ When cells have high Δψm, JC-1 forms J-aggregates in mitochondria and exhibits red fluorescence. However, if Δψm decreases due to the damage of mitochondrial membrane, JC-1 leaks into the cytoplasm and shifts to J-monomers, which emits green fluorescence. Therefore, red/green ratio of cell population with JC-1 staining correlates with Δψm and represents mitochondrial membrane integrity. ^43^

Interestingly, SAMHD1 expression decreased Δψm in the absence or presence of HIV-1-Luc/VSV-G infection although infection potentiated the reduction (Figures 4A and S4A). At 2, 3 and 4 dpi, HIV-1-Luc/VSV-G infection significantly decreased Δψm of THP-1 Ctrl cells, while the reduction in SAMHD1 KO cells was milder compared with that in THP-1 Ctrl cells. NVP treatment partially reverted Δψm reduction in both infected THP-1 Ctrl cells and SAMHD1 KO cells despite complete inhibition of HIV-1-Luc/VSV-G replication (Figures 4A vs. 1A). This was consistent with the observation of partial inhibition of HIV-1-Luc/VSV-G induced apoptosis by NVP treatment (Figure 1). Furthermore, SAMHD1 expression was reconstituted in SAMHD1 KO cells to confirm the role of SAMHD1 in mitochondrial depolarization with or without HIV-1-Luc/VSV-G infection. At 2 and 3 dpi, THP-1 Ctrl and SAMHD1 KI cells showed significantly less Δψm, which was further decreased by HIV-1 infection, compared to SAMHD1 KO and Lvx cells (Figures 4B and S4B). These results indicated that SAMHD1 depolarized mitochondria, which was enhanced upon HIV-1 infection.

**Figure 4.**
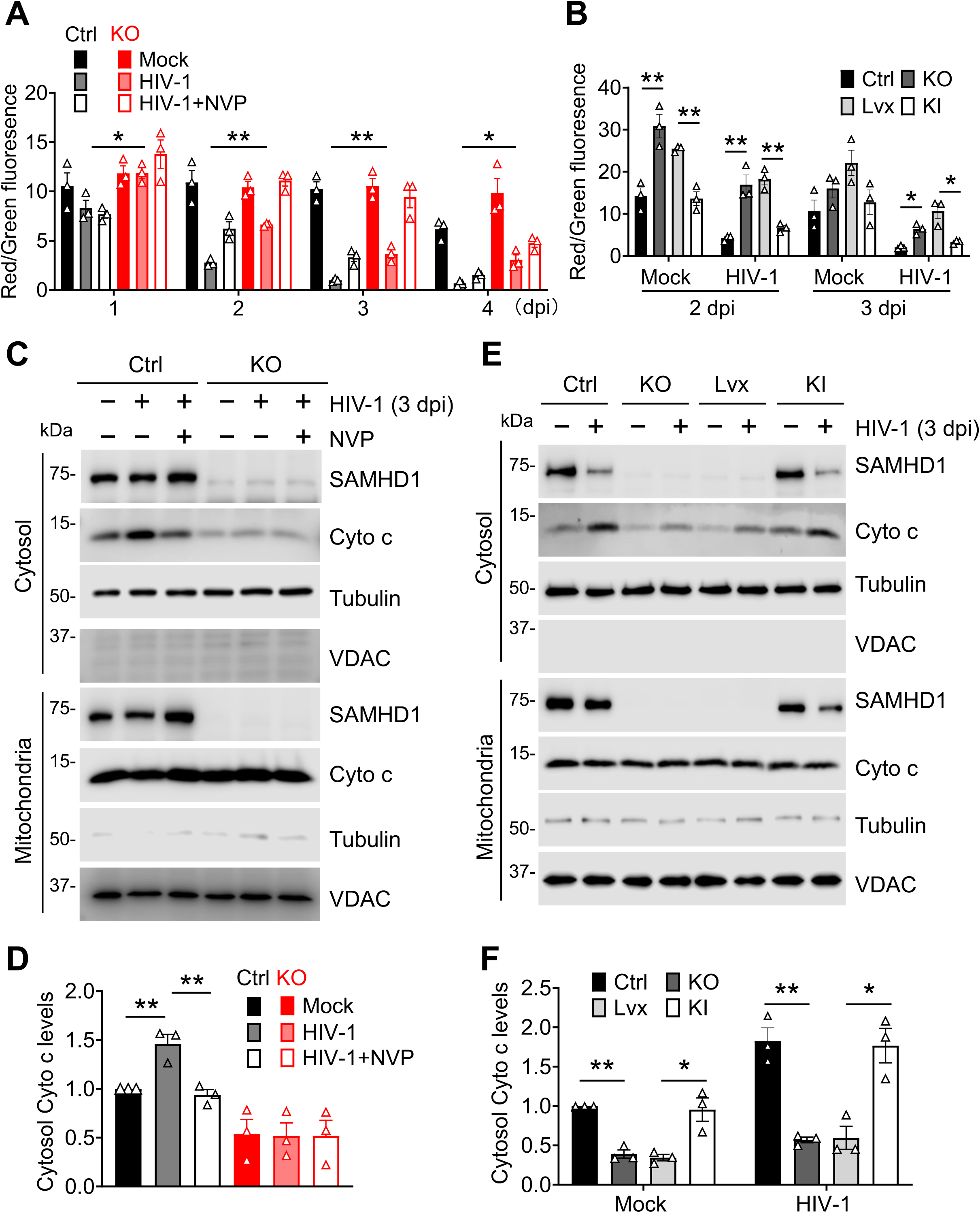
SAMHD1 enhances apoptosis induced by HIV-1 infection through the mitochondrial pathway. **(A, C, and D)** THP-1 Ctrl and SAMHD1 KO cell lines were infected with HIV-1-Luc/VSV-G (MOI=2) or mock-treated for the indicated times. NVP was used to block HIV-1 infection. **(B, E, and F)** THP-1 Ctrl, SAMHD1 KO, Lvx control, and SAMHD1 KI cell lines were infected with HIV-1-Luc/VSV-G (MOI=2) for the indicated times. **(A and B)** Cells were stained with JC-1 and measured by flow cytometry. Original flow cytometry results are shown in Fig. S4A and S4B, respectively. **(C and E)** Detection of SAMHD1, cytochrome c (Cyto c), VDAC, and tubulin in the cytosol and mitochondrial fractions at 3 dpi by Western blot. VDAC and tubulin were used as mitochondrial and cytosolic markers, respectively. **(D and F)** The relative levels Cyto c in the cytosol fraction at 3 dpi (C and E, respectively) were quantified by densitometry analysis and normalized to tubulin. The level of Cyto c in the cytosol fraction of mock-infected THP-1 Ctrl cells was set to 1. **(A, B, D, and F)** The unpaired t-test was used for statistical significance. *p < 0.05; **p < 0.01.

Depolarized mitochondria release Cyto c to initiate mitochondrial apoptotic pathway. ^44^ We therefore examine whether SAMHD1 expression also promoted cytosolic release of Cyto c. Subcellular fractionation was performed in THP-1 Ctrl and SAMHD1 KO cells mock infected or infected with HIV-1-Luc/VSV-G and treated with NVP for 3 dpi to detect cytosolic Cyto c. The cytosolic and mitochondrial fractions of cell lysate were detected by Western blot. Tubulin was used as the cytosolic marker and voltage-dependent anion channel (VDAC) was used as the mitochondrial marker. SAMHD1 was detected in both the cytosolic and mitochondrial fractions. Cyto c was more enriched in the cytosolic fraction in THP-1 Ctrl cells than SAMHD1 KO cells, suggesting that SAMHD1 expression enhanced cytosolic release of Cyto c (Figures 4C and 4D). HIV-1-Luc/VSV-G infection further promoted cytosolic Cyto c in THP-1 Ctrl cells, which was mitigated by NVP treatment.

HIV-1-Luc/VSV-G induced cytosolic release of Cyto c was alleviated in SAMHD1 KO cells, suggesting that HIV-1-induced cytosolic release of Cyto c depends on SAMHD1 expression. SAMHD1 expression was reconstituted in SAMHD1 KO cells to confirm the role of SAMHD1 in cytosolic release of Cyto C. THP-1 Ctrl, SAMHD1 KO, Lvx and SAMHD1 KI cell lines were infected with HIV-1-Luc/VSV-G or mock infected for 3 dpi. Consistently, we found that reconstitution of SAMHD1 expression in SAMHD1 KO cells reinforced cytosolic release of Cyto c in the absence or presence of HIV-1-Luc/VSV-G infection to the level comparable to THP-1 Ctrl cells (Figures 4E and 4F). Together, these data suggest that SAMHD1 enhances apoptosis induced by HIV-1 infection through the mitochondrial pathway by depolarizing mitochondria and cytosolic release of Cyto c.

### SAMHD1-enhanced apoptosis is associated with increased BIK expression independently of HIV-1 infection

BCL-2 family proteins play an important role in the regulation of the mitochondrial pathway. ^27,28^ To investigate the mechanism by which SAMHD1 enhances apoptosis induced by HIV-1 infection through the mitochondrial pathway, the expressions of BCL-2 family proteins in THP-1 Ctrl and SAMHD1 KO cell lines infected with HIV-1_NL4-3_ and with or without NVP at 2 to 8 dpi were measured by Western blot. The expression of BIK in THP-1 Ctrl cells, with or without HIV-1 infection, was higher than that in SAMHD1 KO cells at 2, 4, 6 and 8 dpi (Figures 5A, 5B and S5A), while the expressions of BCL-2, BCL-X_L_, BAX, BAK, BIM and BID were not affected by SAMHD1 expression, HIV-1 infection, or NVP treatment (Figure S5A), suggesting that SAMHD1-enhanced apoptosis is associated with increased BIK expression. Interestingly, BIK expression was increased by HIV-1 infection at 8 dpi only in THP-1 Ctrl cells but not in SAMHD1 KO cells based on the average of 3 independent experiments (Figure 5B).

**Figure 5.**
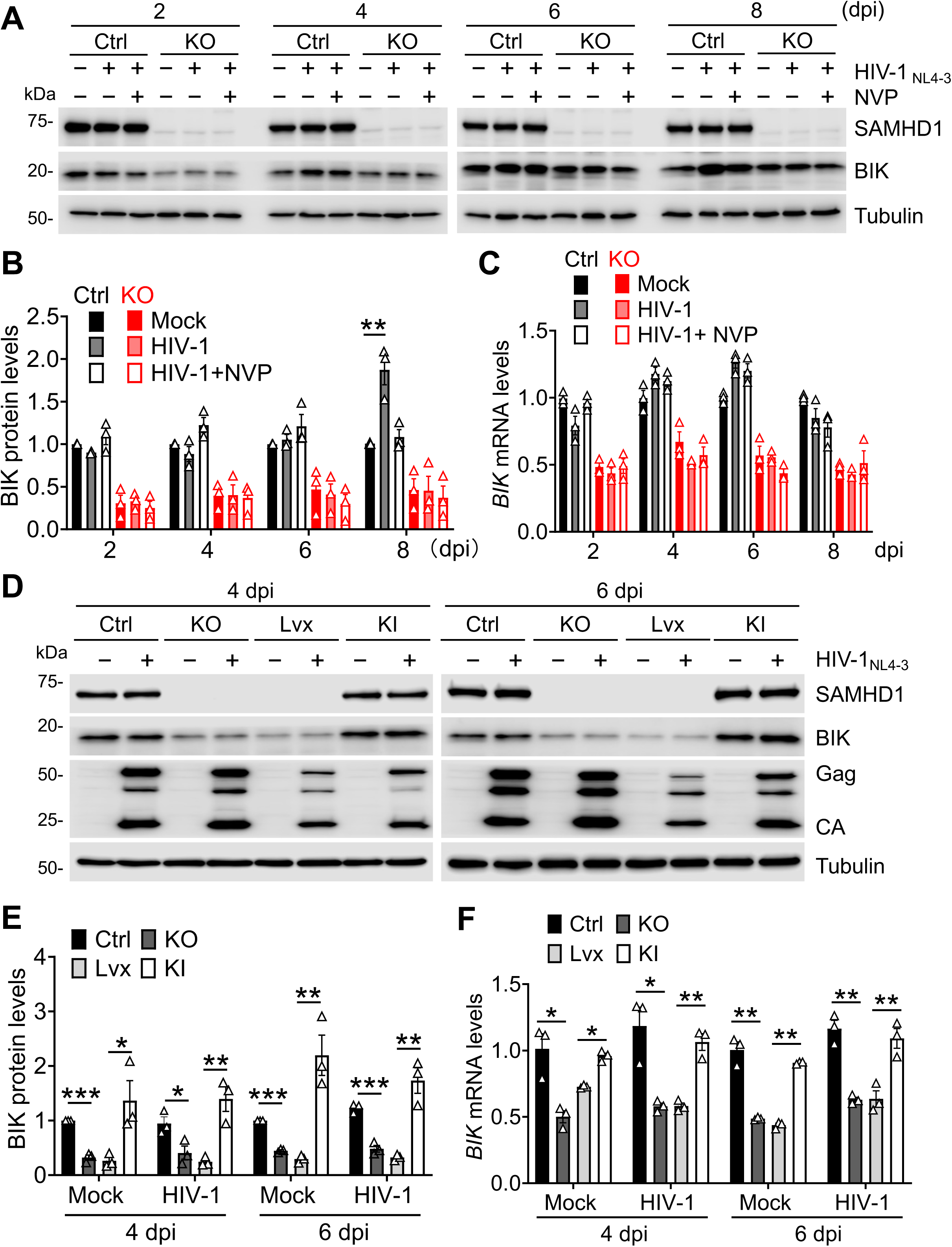
SAMHD1-enhanced apoptosis is associated with increased BIK expression independently of HIV-1 infection. **(A, B, and C)** THP-1 Ctrl and SAMHD1 KO cell lines were infected with HIV-1_NL4-3_ (MOI=2) for indicated time or mock-treated. NVP was used to block HIV-1 infection. **(D, E, and F)** THP-1 Ctrl, SAMHD1 KO, Lvx control, and SAMHD1 KI cell lines were infected with HIV-1_NL4-3_ (MOI=2) or mock-treated for the indicated time. **(A and D)** SAMHD1, BIK, tubulin, and HIV-1 Gag and CA in cells were detected by Western blot. **(B and E)** The relative levels of BIK protein were quantified by densitometry analysis and normalized to tubulin. BIK protein levels of mock-infected THP-1 Ctrl cells were set to 1. **(C and F)** The mRNA levels of BIK were detected by qRT-PCR and were expressed relative to mock-infected THP-1 Ctrl cells that was set to 1. GAPDH was used to normalize qRT-PCR results. **(B, C, E, and F)** The unpaired t-test was used for statistical significance. *p < 0.05; **p < 0.01; ***p < 0.001. The data (means ± SEM) represent three independent experiments.

To investigate whether SAMHD1 affects BIK expression through proteasomal or lysosomal degradation, THP-1 Ctrl and SAMHD1 KO cell lines were treated separately with proteasomal or lysosomal inhibitors 1-2 days before BIK and SAMHD1 detection by Western blot. MG132 is a proteasome inhibitor that induces PARP cleavage,^45–47^ while Chloroquine (CQ) is a lysosomal hydrolase inhibitor that increases the autophagy marker light chain 3 (LC3) isoform LC3A/B-II accumulation.^48,49^ Cleaved PARP and LC3A/B-I/II accumulation were used as positive controls, confirming the validity of the MG132 and CQ treatments. BIK expression was not affected by MG132 or CQ treatment (Figure S6A), suggesting that SAMHD1 does not regulate BIK expression through the proteasomal or lysosomal degradation pathway.

Our previous study^18^ and this study showed that SAMHD1 is localized at the mitochondria and cytosol (Figures 4C and 4E). BIK is localized in the endoplasmic reticulum (ER) and translocated to mitochondria when the mitochondrial apoptotic pathway is activated.^50^ To investigate whether SAMHD1 is associated with BIK, we performed co-immunoprecipitation (Co-IP) in THP-1 Ctrl cells and SAMHD1 KO cells and we found that SAMHD1 did not interact with BIK in THP-1 cells (Figure S6B). Then, we further investigated whether HIV-1_NL4-3_ infection affected the interaction between SAMHD1 and BIK in THP-1 Ctrl cells using Co-IP. No interaction between SAMHD1 and BIK was observed in THP-1 cells regardless of HIV-1_NL4-3_ infection (Figure S6C).

To assess whether BIK expression was affected by SAMHD1 at the transcription level, THP-1 Ctrl and SAMHD1 KO cells were mock infected or infected with HIV-1_NL4-3_ and treated with NVP. At 2 to 8 dpi, BIK mRNA was measured by qRT-PCR. Overall, the *BIK* mRNA levels were higher in THP-1 Ctrl cells compared to SAMHD1 KO cells, regardless of HIV-1_NL4-3_ infection or NVP treatment (Figure 5C), suggesting that SAMHD1 increases BIK expression at the mRNA level. *BAX* mRNA, used as a negative control, was not affected with SAMHD1 expression (Figure S5C).

Next, SAMHD1 was reconstituted in SAMHD1 KO cells to confirm its effect on upregulation of BIK expression. At 4 and 6 dpi, we observed that the levels of BIK protein (Figures 5D and 5E) and mRNA (Figure 5F) in THP-1 Ctrl and SAMHD1 KI cells were higher than these in SAMHD1 KO and Lvx cells, regardless of HIV-1_NL4-3_ infection, suggesting that SAMHD1 increases BIK expression at both the protein and mRNA levels. In contrast, the protein levels of BCL-2, BCl-X_L_, BAX, BAK, BIM and BID (Figure S5B), as well as mRNA levels of *BAX* (Figure S5D), were not affected by SAMHD1 expression or HIV-1_NL4-3_ infection of THP-1 cell lines. These data indicate that SAMHD1-enhanced apoptosis is associated with increased BIK expression, independently of HIV-1 infection.

### Endogenous BIK contributes to SAMHD1-enhanced apoptosis induced by HIV-1 infection

To investigate the role of BIK in SAMHD1-enhanced apoptosis, we knocked out BIK expression in THP-1 SAMHD1 Ctrl and SAMHD1 KO cell lines and then measured HIV-1 infection and apoptosis (Figures 6A and 6B). BIK knockout was confirmed with Western blot (Figure 6C). We established six stable THP-1 cell lines with Ctrl empty guide RNA vector (V), Ctrl BIK KO-2 (BIK gRNA-2), Ctrl BIK KO-4 (BIK gRNA-4), SAMHD1 KO V, SAMHD1 and BIK double-KO (DKO)-2 (BIK gRNA-2), and SAMHD1 and BIK DKO-4 (BIK gRNA-4). These cell lines were infected with HIV-1-Luc/VSV-G or mock infected. At 3 dpi, HIV-1-Luc/VSV-G infection in THP-1 cells was not significantly affected when BIK was depleted (Figure 6A). The total percentages of apoptosis in HIV-1 infected THP-1 Ctrl V cells were significantly higher than these observed in Ctrl BIK KO-2 and Ctrl BIK KO-4 cells (Figures 6B and S7A), suggesting that BIK depletion decreased HIV-1-induced apoptosis in THP-1 SAMHD1 Ctrl cells. In contrast, BIK depletion had no effect on HIV-1-induced apoptosis in SAMHD1 KO cells (Figures 6B and S7A). We further investigated the role of BIK in SAMHD1-enhanced apoptosis by analyzing the cleavage of caspases 3/7 and PARP. BIK depletion reduced HIV-1-induced caspases 3/7 and PARP cleavage in THP-1 Ctrl cells. Comparatively, loss of BIK did not significantly reduce HIV-1-induced caspases 3/7 and PARP cleavage in SAMHD1 KO cell lines (Figures 6C and 6D).

**Figure 6.**
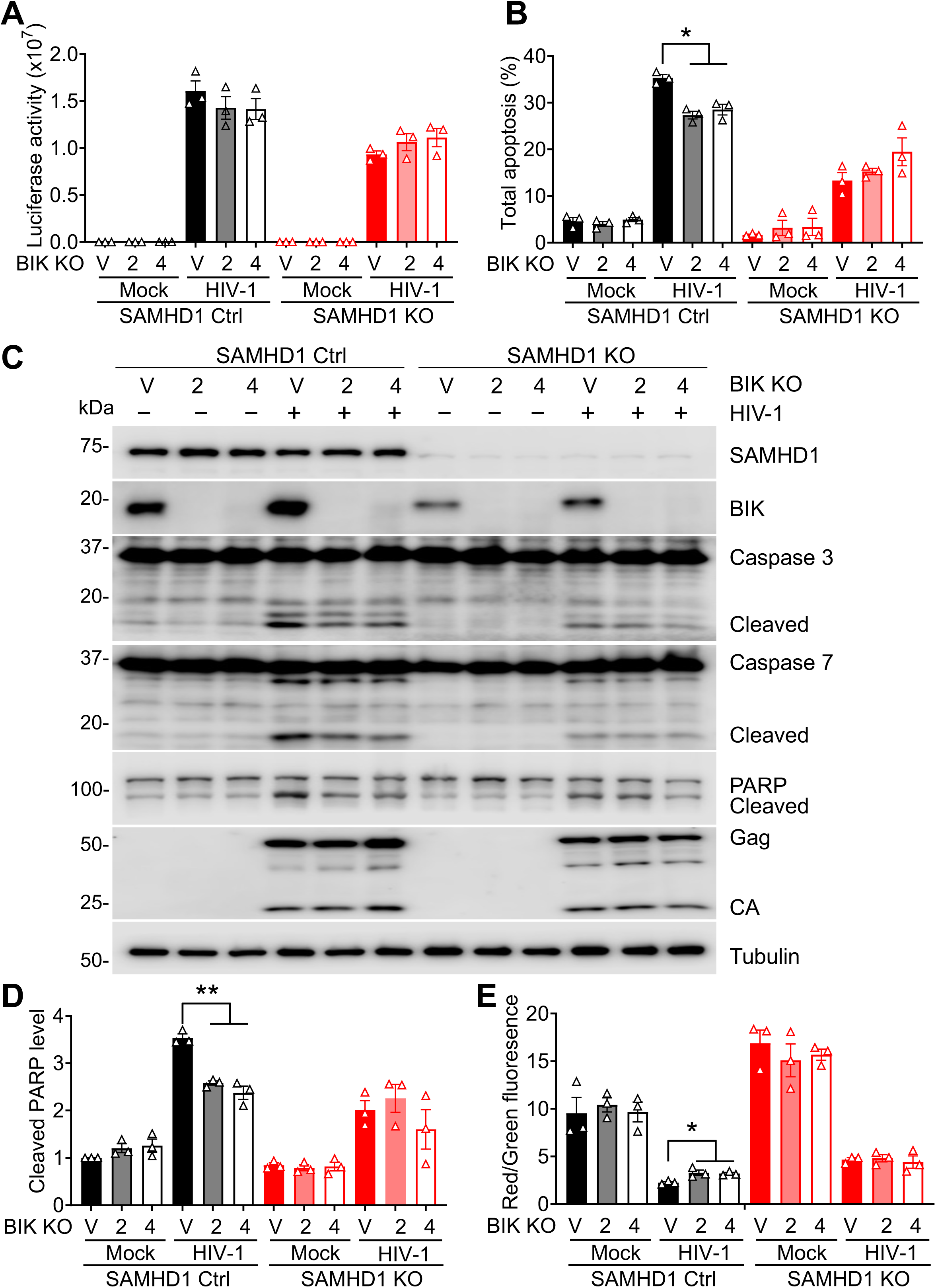
Endogenous BIK contributes to SAMHD1-enhanced apoptosis induced by HIV-1 infection. **(A-E)** THP-1 Ctrl and SAMHD1 KO cell lines expressing empty guide RNA vector (V), BIK-specific guide RNA 2 (BIK KO-2), BIK-specific guide RNA 4 (BIK KO-4) were infected with HIV-1-Luc/VSV-G (MOI=2) or mock infected. Cells were harvested at 3 dpi for further analyses. **(A)** HIV-1 infection levels were measured by a luciferase assay and normalized for cellular protein concentration. **(B)** Total apoptosis was measured by flow cytometry as indicated in Fig. 1B. Original flow cytometry results are shown in Fig. S7A. **(C)** Detection of SAMHD1, BIK, caspase 3, caspase 7, PARP, HIV-1 Gag and CA, and tubulin by Western blot. Tubulin was used as a loading control. **(D)** The relative cleaved PARP levels were quantified by densitometry analysis and normalized to tubulin. The level of cleaved PARP in mock-infected THP-1 Ctrl EV cells was set to 1. **(E)** Cells were stained by JC-1 and measured by flow cytometry. Original flow cytometry results are shown in Fig. S7B. The data in A, B, D and E (means ± SEM) represent three independent experiments. The unpaired t-test was used for statistical significance.

BIK is a BH3-only protein that converges apoptotic stimuli, permeabilizes outer mitochondrial membrane by activating BAX/BAK oligomerization through releasing calcium ions from ER and initiates mitochondrial apoptosis.^50^ We therefore questioned whether BIK was involved in SAMHD1-mediated mitochondrial depolarization. THP-1 Ctrl or SAMHD1 KO cell lines with or without BIK depletion were infected with HIV-1-Luc/VSV-G or mock infected. At 3 dpi, the red/green fluorescence ratio of JC-1 dye staining in THP-1 Ctrl V cells infected with HIV-1 was significantly lower compared to Ctrl BIK KO-2 and Ctrl BIK KO-4 cells, suggesting that BIK depletion replenished Δψm in HIV-1-infected THP-1 Ctrl cells. Conversely, BIK depletion did not further promote Δψm of HIV-1-infected SAMHD1 KO cells (Figures 6E and S7B). These data indicate that endogenous BIK mediates SAMHD1-induced mitochondrial depolarization in THP-1 cells, suggesting that BIK contributes to SAMHD1-enhanced mitochondrial apoptosis induced by HIV-1 infection.

## DISCUSSION

Apoptosis is a major form of programmed cell death and plays a crucial role in various physiological processes, including viral infections.^27,28^ In this study, we identified that SAMHD1 enhances apoptosis induced by HIV-1 infection in THP-1 cells through the mitochondrial pathway (summarized in Figure 7). Other studies have shown that SAMHD1 enhances apoptosis in primary human monocytic cells infected with HTLV-1^39^ and overexpression of SAMHD1 in non-small cell lung cancer A549 cells induces apoptosis.^51^ Conversely, knock-down SAMHD1 increases apoptosis in ovarian cancer cell lines.^52^ SAMHD1 silencing also enhances apoptotic cell death in lung adenocarcinoma cells. ^53^ Moreover, SAMHD1 downregulation combined with radiation enhance anti-tumor immune responses to lung adenocarcinoma in mice.^53^ These studies suggest that SAMHD1-mediated apoptotic cell death may be dependent on the cell types and apoptotic stimuli, which could be used to develop potential therapeutic strategies against cancers.

**Figure 7.**
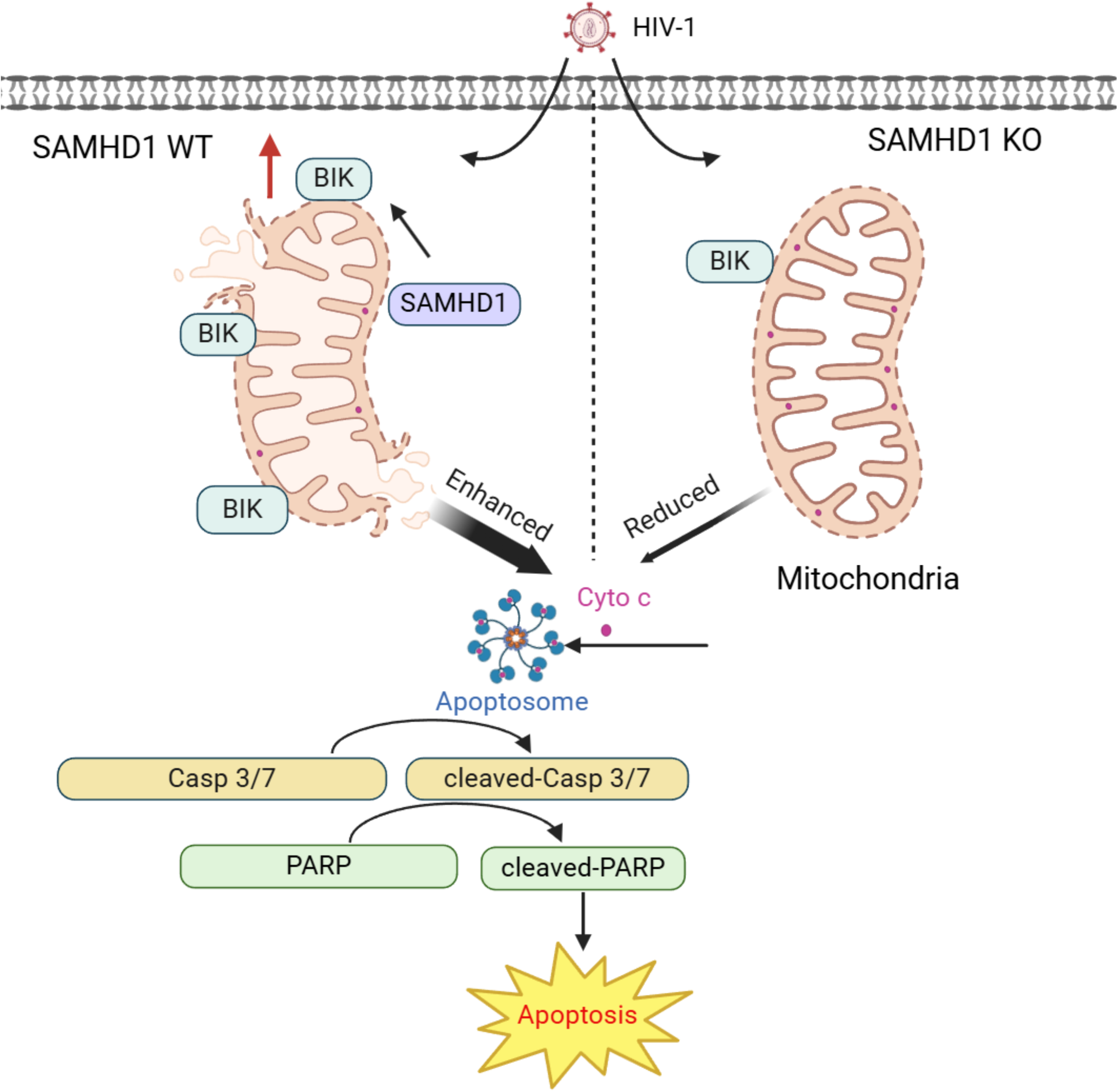
SAMHD1 enhances HIV-1-induced apoptosis in monocytic cells via the mitochondrial pathway. Endogenous SAMHD1 enhances HIV-1-induced apoptosis in undifferentiated THP-1 cells. Mechanistically, SAMHD1 increases mitochondrial membrane damage, BIK expression, and Cyto c release into the cytosol of HIV-1 infected cells. As a result, caspases 3/7 are activated by apoptosomes and then cleave downstream PARP, leading to enhanced apoptosis in the presence of SAMHD1. In contrast, knockout of endogenous SAMHD1 reduces apoptosis in HIV-1-infected THP-1 cells. Our findings suggest positive regulatory mechanisms by which SAMHD1 enhances apoptosis induced by HIV-1 infection through the mitochondrial pathway.

SAMHD1 did not inhibit HIV-1 infection at 2 and 3 dpi in dividing THP-1 cells (Figure 1A). In our previous study, SAMHD1 inhibits HIV-1 infection at 1 and 2 dpi in THP-1 cell-differentiated macrophages-like cells with increased SAMHD1 protein expression.^24,54^ Lower dNTP pool in nondividing macrophages-like cells (higher SAMHD1 expression) does not support the HIV-1 reverse transcription, whereas a higher concentration of dNTPs in proliferating monocytic cells (lower SAMHD1 expression) reaches the threshold required for efficient reverse transcription.^24,54^ Treatment of NVP partially inhibited HIV-1-induced apoptosis or the reduction of Δψm (Figures 1 and 4). Single-cycle HIV-1-Luc/VSV-G contains Vpr, a viral protein known to induce apoptosis.^37,38^ NVP treatment of cells inhibits HIV-1 reverse transcription but does not affect Vpr incorporated in virions and its expression during infection, which can lead to apoptosis.

In this study, we observed that SAMHD1 enhances HIV-1-induced apoptosis in THP-1 cells by reducing Δψm. A previous study showed that SAMHD1 knockout impairs Δψm in mouse macrophages at both resting state and M1 polarized state.^55^ Thus, the function of SAMHD1 in maintaining Δψm varies across different cell types. Our previous study showed that SAMHD1 enhances Fas-L induced apoptosis through the death receptor apoptotic pathway. ^40^ In our study, truncated BID (tBID), which is cleaved from BID by caspase 8 in the death receptor apoptotic pathway,^56,57^ was not detected in monocytic THP-1 cells during HIV-1 infection (Figures S5A and S5B). This indicates that SAMHD1-enhanced apoptosis in HIV-1-infected cells does not occur through the death receptor apoptotic pathway. Therefore, SAMHD1 can enhance both the death receptor and mitochondrial apoptotic pathways, depending on the specific stimuli.

In this study, we observed that SAMHD1-enhanced apoptosis is associated with increased BIK expression. BIK is a pro-apoptotic member of the BCL-2 family and binds to anti-apoptotic proteins BCL-2 and BCL-X_L_. The interaction between BIK and BCL-2 or BCL-X_L_ inhibits the function of anti-apoptotic proteins, leading to induces MOMP and Cyto c release^28,50^. A previous study showed that knockdown of BIK in human airway epithelial cells significantly reduced the apoptosis induced by influenza A virus infection, suggesting that BIK is a major mediator for the apoptotic pathway triggered by viral infection.^58^ Our previous^18^ and this study (Figures 4C and 4E) showed that SAMHD1 is localized at mitochondria, as confirmed by cytosolic and mitochondrial fractionation. However, we did not detect the interaction between endogenous SAMHD1 and BIK in THP-1 cells. The loss of BIK expression selectively reduces the enhancement of SAMHD1 in HIV-1-induced apoptosis in THP-1 Ctrl cells but does not influence apoptosis levels in SAMHD1 KO cells (Figure 6). The phenomenon occurs likely because expression of BIK in THP-1 Ctrl cells is higher than in SAMHD1 KO cells (Figure 5). Therefore, the loss of higher BIK expression in Ctrl cells leads to a more pronounced reduction in apoptosis compared to SAMHD1 KO cells.

The deprived BIK protein level in SAMHD1 KO cells might have failed the required threshold for BIK protein to trigger apoptosis for HIV-1 infection in THP-1 cells (Figure 6). Future investigation is required to confirm this possibility. Additional experiments can be performed with reconstitution of BIK expression, use of BIK-specific apoptotic stimuli, and use of other cell types and primary monocytes having different levels of SAMHD1 or BIK expression. Alternatively, BIK protein might work with SAMHD1 protein to execute HIV-1 induced apoptosis. Loss of BIK expression reverted Δψm in SAMHD1 expressing THP-1 cells, but not in SAMHD1 KO cells, upon HIV-1 infection, suggesting that BIK is required for SAMHD1-enhnaced mitochondrial depolarization by HIV-1 (Figures 6E and S7B). HIV-1 infection to SAMHD1 expressing THP-1 cells did not further promote BIK protein expression by 4 dpi at which SAMHD1-enhanced apoptosis was already observed (Figures 5A and 5B vs. 3A and 3B). BIK and SAMHD1 interaction in the absence or presence of HIV-1 infection was not detected (Figure S6C). Therefore, other cellular factors and mechanisms might be involved in SAMHD1/BIK dependent mitochondrial apoptosis induced by HIV-1 infection. A prior study showed that SAMHD1 interacts with the mitochondrial protein VDAC1 in mouse macrophages.^55^ We have reported that SAMHD1 interacts with the mitochondrial antiviral-signaling protein in THP-1 cells ^18^. It is also possible that SAMHD1 interactions with other mitochondrial proteins contribute to enhanced mitochondrial apoptosis.

The dNTP hydrolase (dNTPase) activity of SAMHD1 is critical for its HIV-1 restriction function in non-dividing immune cells.^4,6,54,59–61^ We have reported that the dNTPase activity of SAMHD1 is important for its suppression of innate immune responses in differentiated monocytic cells, but not in dividing HEK293T cells.^17,18,62^ Further study is required to investigate whether dNTPase activity of SAMHD1 is important for its function in enhancing HIV-1-induced mitochondrial apoptosis. In this type of future study, it is challenging but necessary to distinguish SAMHD1-mediated HIV-1 restriction from the effects on virus-induced apoptosis in cells.

We observed that SAMHD1 enhanced the MG-132 induced apoptosis in THP-1 cells (Figure S6A), suggesting that SAMHD1 plays an important role in regulating cell death. An *in vivo* study using humanized mice and nonhuman primates reported that treatment of lung CD4+ T cells with the pan-caspase inhibitor Z-VAD-FMK and caspase 3 inhibitor Z-DEVD-FMK decrease cell death induced by HIV-1 and SIV infection, suggesting HIV-1-induced apoptosis significantly contributes to the loss of CD4+ T cells.^63^ Further studies are needed to investigate the role of SAMHD1 in HIV-1-induced loss of CD4+ T cells.

In summary, we demonstrated that endogenous SAMHD1 enhances apoptosis induced by HIV-1 infection through the mitochondrial pathway in monocytic cells. Our results provide new insights into the mechanisms underlying the apoptotic function of SAMHD1 during HIV-1 infection.

## ACKNOWLEDGMENTS

We thank the Wu lab members for helpful discussions and suggestions. We appreciate the reagents provided by the National Institutes of Health (NIH) AIDS Reagent Program. Research reported in this publication was supported by the National Institute of Allergy and Infectious Diseases of the NIH under Award Number R01AI141495. L. W. is also supported by NIH grants R61AI169659, R21AI170070, R21AI181742, and P30CA086862-24S1. The content is solely the responsibility of the authors and does not necessarily represent the official views of the NIH.

## AUTHOR CONTRIBUTIONS

Conceptualization, H.Y. and L.W.; methodology, H.Y.; validation, H.Y.; formal analysis, H.Y.; investigation, H.Y.; resources, L.W.; visualization, H.Y. and L.W.; funding acquisition, L.W.; writing– original draft, H.Y., P.H.C. and L.W.; writing– review & editing, H.Y., P.H.C. and L.W.

## SUPPLEMENTAL INFORMATION

Document S1. Figures S1–S7

## METHODS

### KEY RESOURCES TABLE

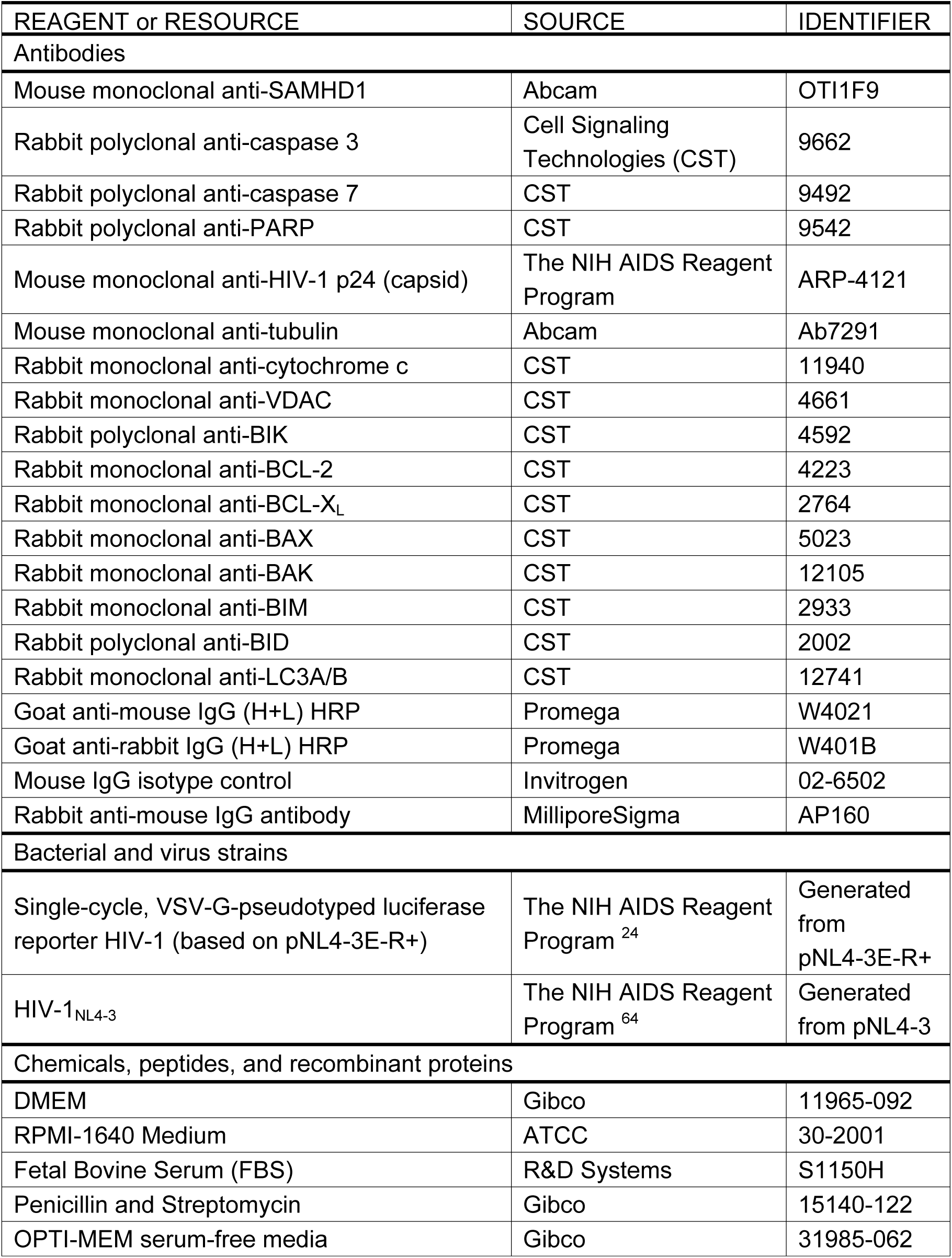
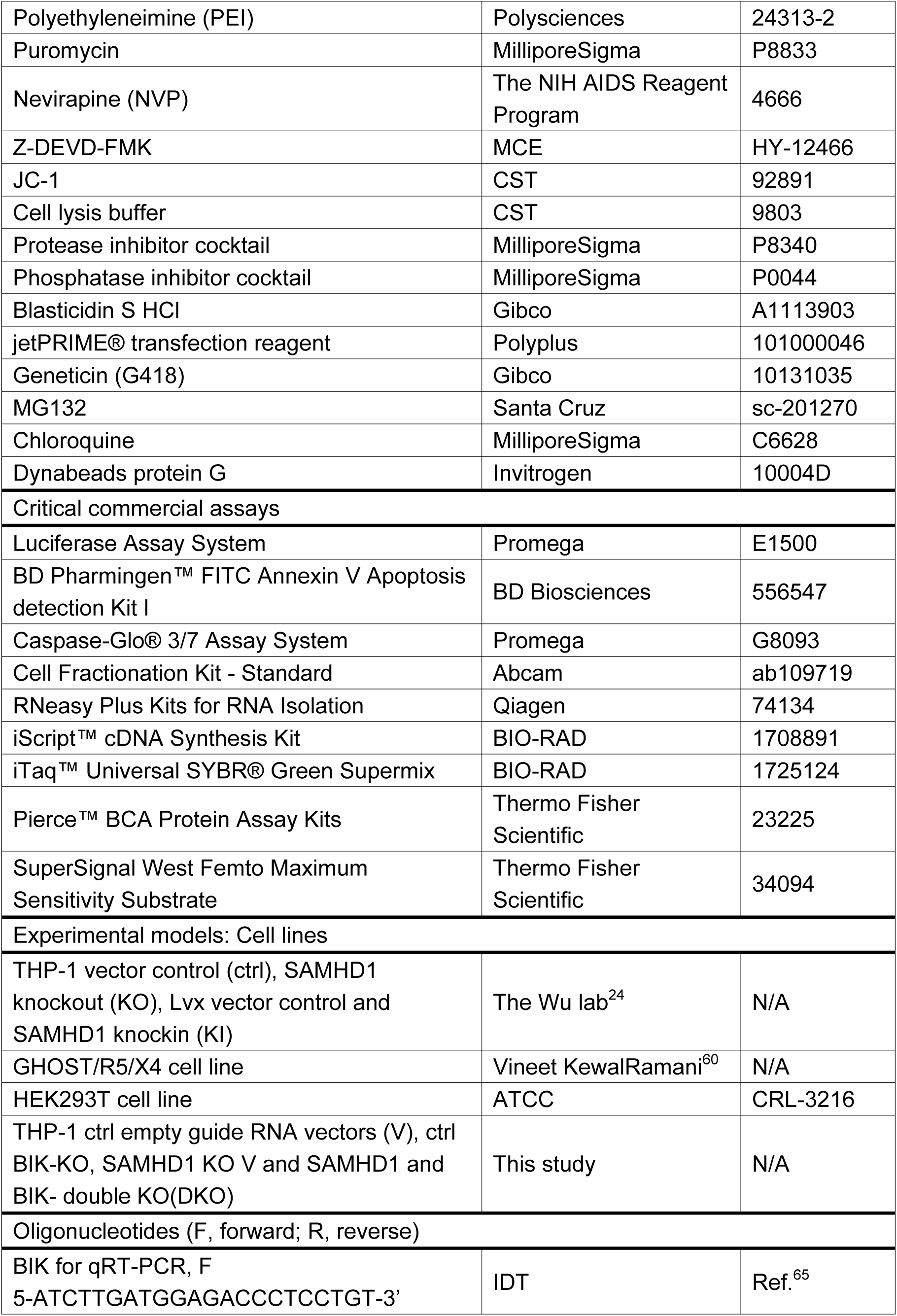
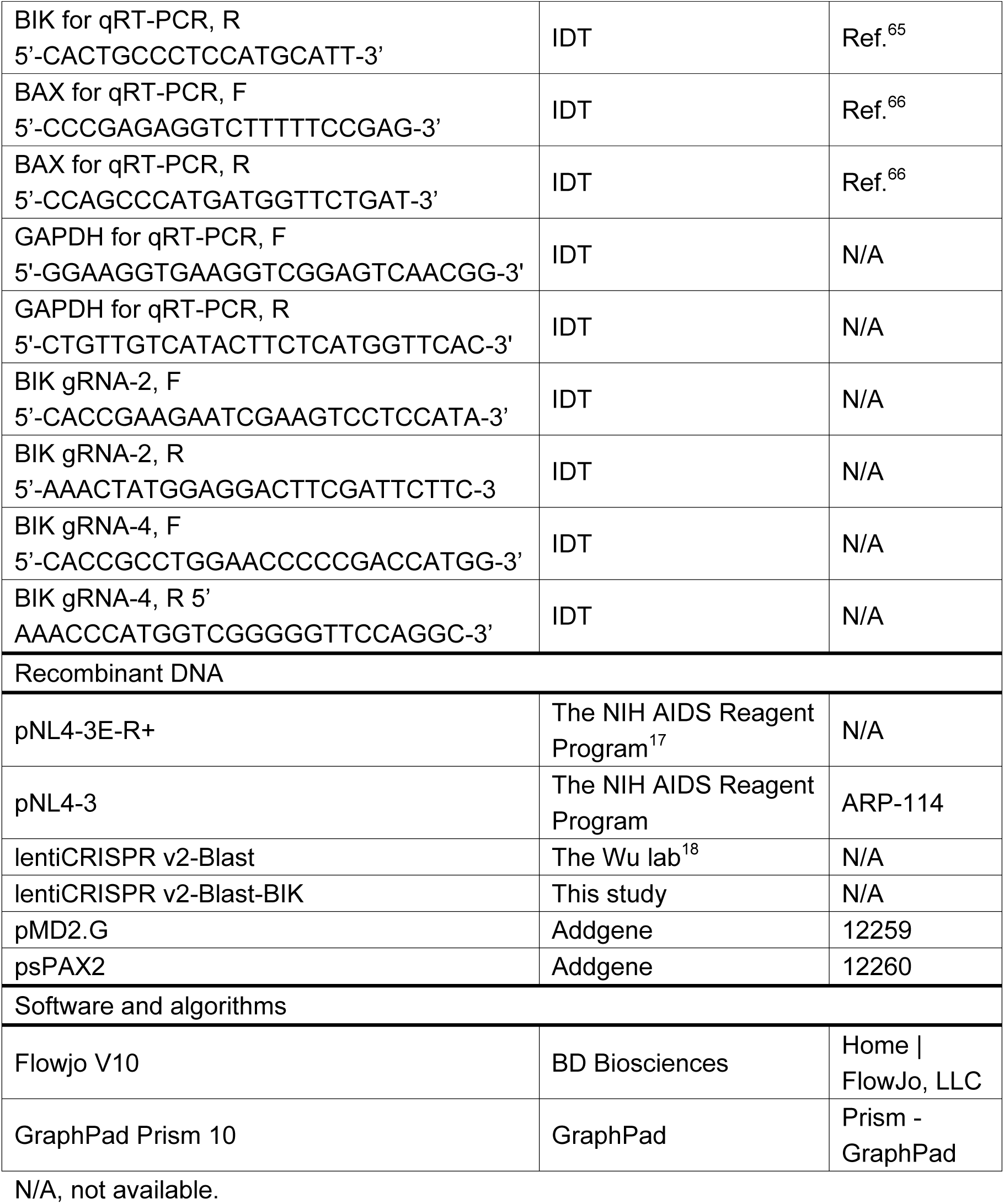

### RESOURCE AVAILABILITY

#### Lead contact

Further information and requests for resources and reagents should be directed to and will be fulfilled by the lead contact, Li Wu.

#### Materials availability

Unique reagents generated in this study and listed in above table can be requested from the lead contact, Li Wu.

### METHODS DETAILS

#### Cell lines

THP-1 Ctrl, SAMHD1 KO, Lvx and SAMHD1 KI cell lines were described previously.^17,24^ HEK293T cells and GHOST/R5/X4 cells were described previously.^17^ HEK293T were cultured in DMEM media with 10% FBS, 100 U/mL penicillin, and 100 μg/mL streptomycin. GHOST/R5/X4 were cultured in DMEM media with 10% FBS, 100 U/mL penicillin, 100 μg/mL streptomycin, 500 μg/mL neomycin, 50 μg/mL hygromycin B and 1 μg/mL puromycin. THP-1 BIK KO cell lines were generated by CRISPR/Cas9 technology and grown in RPMI 1640 media with 10% fetal bovine serum (FBS), 100 U/mL penicillin, 100 μg/mL streptomycin, and 1 μg/mL puromycin and 10 μg/mL blasticidin.

#### CRISPR/Cas9-mediated gene knockout

HEK293T were transfected with lentiCRISPR v2-Blast, or lentiCRISPR v2-Blast-BIK gRNA, psPAX2 and pMD2.G (ratio 4: 3 :1) by using jetPRIME® transfection reagent. Two Guide RNA-target BIK sequences were 5’-TATGGAGGACTTCGATTCTT-3’ and 5’ CCTGGAACCCCCGACCATGG-3’. After 48 h transfection, lentivirus was collected and purified by passing them through a 0.45-micron filter. THP-1 ctrl and SAMHD1 KO cells were transduced with lentivirus in the presence of 10 µg/mL polybrene. At 48 h post-transduction, the cells were cultured in the RPMI 1640 media with the blasticidin (10 μg/mL). The BIK knockout cells were confirmed by Western blot.

#### Virus stocks and viral infection assays

Single-cycle HIV-1-Luc/VSV-G and replication competent HIV-1_NL4-3_ were generated from HEK293T cells. Virus stock was digested with DNase I (40 U/mL) for 1 h at 37 °C and filtered (0.45-micron filter). The infectious units of HIV-1 were measured at limiting dilution on GHOST/X4/R5 cells as described.^54,67^ THP-1 cells were infected with HIV-1-Luc/VSV-G (MOI=2) in the presence of 10 µg/mL polybrene. The infection level of HIV-1 was determined by luciferase assay according to the manufacturer’s instructions. Luciferase values were normalized by protein concentration. THP-1 cells were infected with HIV-1_NL4-3_ (MOI=2) in the presence of 10 µg/mL polybrene by spinoculation at 2,000 g for 2 h at room temperature.

#### Western blot

Western blot analysis was performed as previously described.^19^ Briefly, Cells were lysed in the cell lysis buffer with the protease inhibitor cocktail and phosphatase inhibitor cocktail. The concentration of protein was measured by the bicinchoninic acid (BCA) assay. The same amounts of protein were separated by the SDS-PAGE and transferred onto the membrane. The membranes were blocked in the 5% non-fat milk for 1 h and incubated with indicated primary antibody overnight. The membranes were incubated with HPR-labeled secondary antibody for 1 h and developed by Odyssey Fc Imager. Tubulin was used as a loading control to normalize quantification by densitometry.

#### Cytosolic and mitochondrial fractionation

THP-1 cells were infected with HIV-1-Luc/VSV-G or mock infected for 3 dpi. The cytosolic and mitochondrial fractions of THP-1 cells were separated using Cell Fractionation Kit (Abcam) according to the manufacturer’s instructions. Protease inhibitor cocktail and phosphatase inhibitor cocktail were used in cell lysis buffer. Cytosolic and mitochondrial proteins were analyzed by Western blot. VDAC and tubulin were used as a mitochondrial and cytosolic marker ^18^, respectively.

#### Treatment of cells with NVP, F-DEVD-FMK, MG132, or chloroquine

THP-1 cells were infected with HIV-1-Luc/VSV-G or HIV-1_NL4-3_. NVP (10 mM) or F-DEVD-FMK (20 µM) was maintained in the medium throughout the infection and subsequent culture. THP-1 cells were incubated with MG132 (1 μM) and chloroquine (100 μM) for 3 h and then cultured with fresh medium. After 1 and 2 days of treatment, cells were harvested for Western blot.

#### RNA extraction and qRT-PCR

THP-1 cells were collected after HIV-1 infection for the indicated time. Total RNA was extracted using RNeasy Plus Kits for RNA Isolation according to the manufacturer’s instructions. Total RNA (1 μg per sample) was reverse transcribed using iScript™ cDNA Synthesis Kit. iTaq Universal SYBR Green supermix was used for qPCR. GAPDH mRNA was used as the housekeeping gene to normalize the expression of the target gene^18^.

#### Caspase 3/7 activity assay

Caspase 3/7 activity assay was performed as previously described. ^24^ Briefly, THP-1 cells (1×10^6^ per well) were seeded in the 6-well plates. After the HIV-1 infection for the indicated time, cells (2×10^4^) were transferred into 96-well plates (in 6 replicates). Caspase 3/7 activity was measured by Caspase-Glo® 3/7 Assay System according to the manufacturer’s instructions. Briefly, 100 μL Caspase-Glo® 3/7 Substrate were added into 96-well plates, and the plates were incubated for 1 h at room temperature. The luciferase activity was measured by the microplate Reader (VICTOR Nivo).

#### Co-immunoprecipitation (IP) assay

THP-1 cells were harvested and lysed in the cell lysis buffer. BIK antibody (2 μg per 1×10^7^ cells) or SAMHD1 antibody (2 μg per 1×10^7^ cells) and Dynabeads Protein G magnetic beads were used for IP The same amounts of nonspecific rabbit or mouse IgG were used as a negative control. The bound beads were washed with PBS with 0.1% Tween 20 three times and boiled in protein loading buffer. The input and IP products were detected by Western blot.

#### Statistical analysis

All results were shown as mean ± SEM. GraphPad Prism software was used to analyze data. The unpaired t-test was used for analyses and statistical significance was defined as p < 0.05.

