## Supplement figure S1-S7 for "SAMHD1 enhances HIV-1-induced apoptosis in monocytic cells via the mitochondrial pathway"

**Yang et al. Supplemental figures S1-S7 and legends**

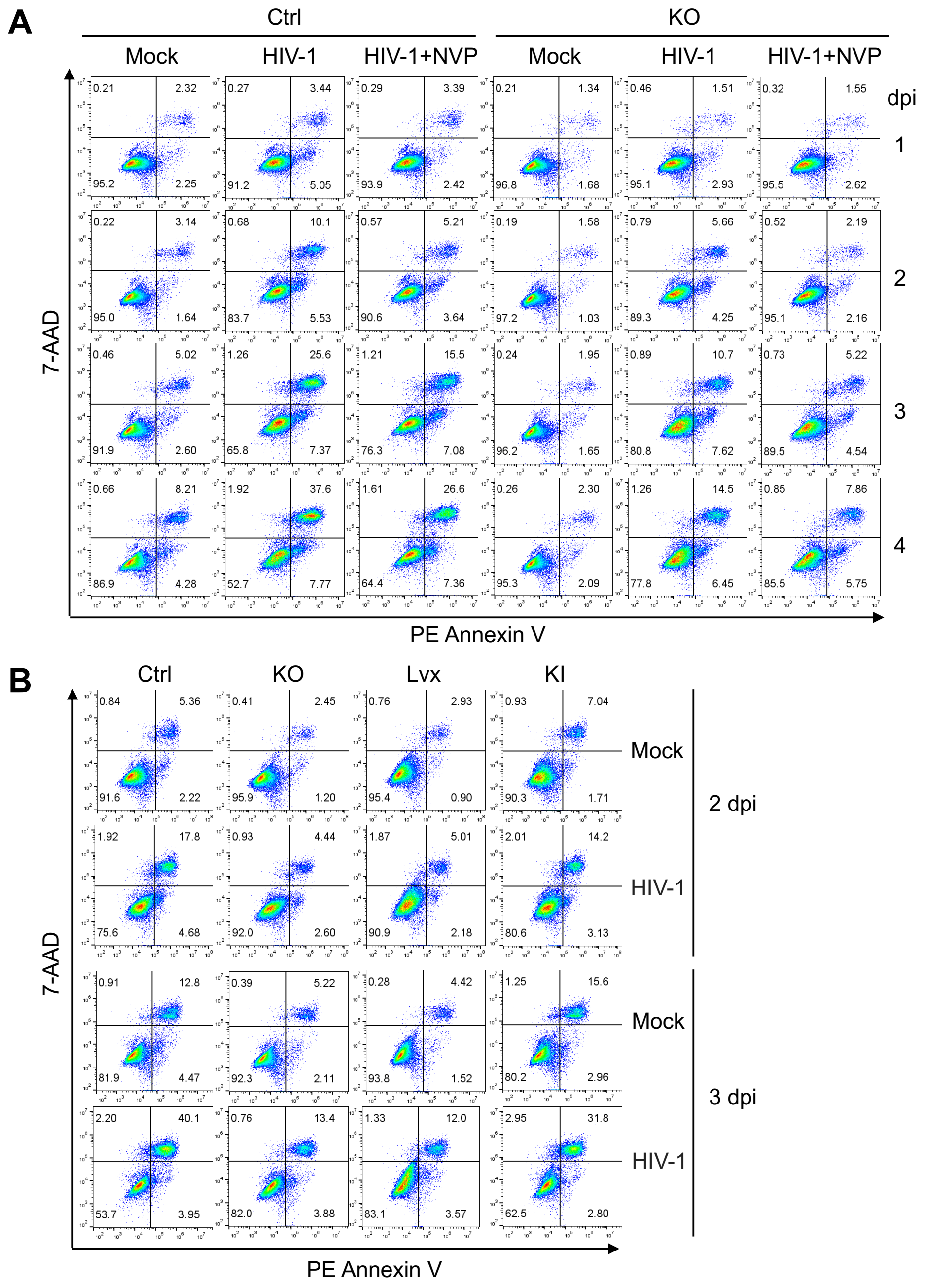


**Figure S1. SAMHD1 enhances apoptosis induced by HIV-1 infection of THP-1 cells**

**(A)** THP-1 control (Ctrl) and SAMHD1 knockout (KO) cells lines were infected with HIV-1-Luc/VSV-G (MOI=2) or mock infected. NVP was used to block viral infection. Cells were harvested at 1-4 days post infection (dpi) for apoptosis analysis.
**(B)** THP-1 Ctrl, SAMHD1 KO, Lvx vector control, and SAMHD1 knock-in (KI) cells lines were infected with HIV-1-Luc/VSV-G (MOI=2) or mock infected. Cells were collected at 2 and 3 dpi for apoptosis analysis.
**(A and B)** Cells were stained with Annexin V and 7-AAD and measured by flow cytometry.

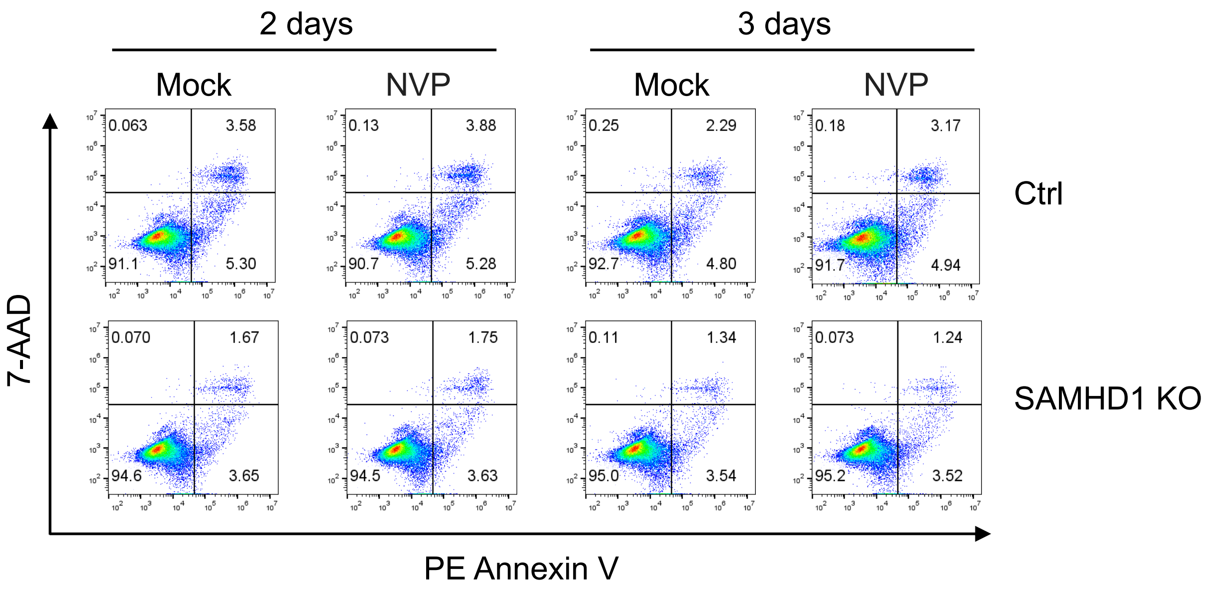


**Figure S2. NVP treatment alone does not affect THP-1 cell apoptosis**

THP-1 Ctrl and SAMHD1 KO cells lines were treated with NVP (10 μM) for 2 or 3 days or mock treated. Cells were stained with Annexin V and 7-AAD and measured by flow cytometry.

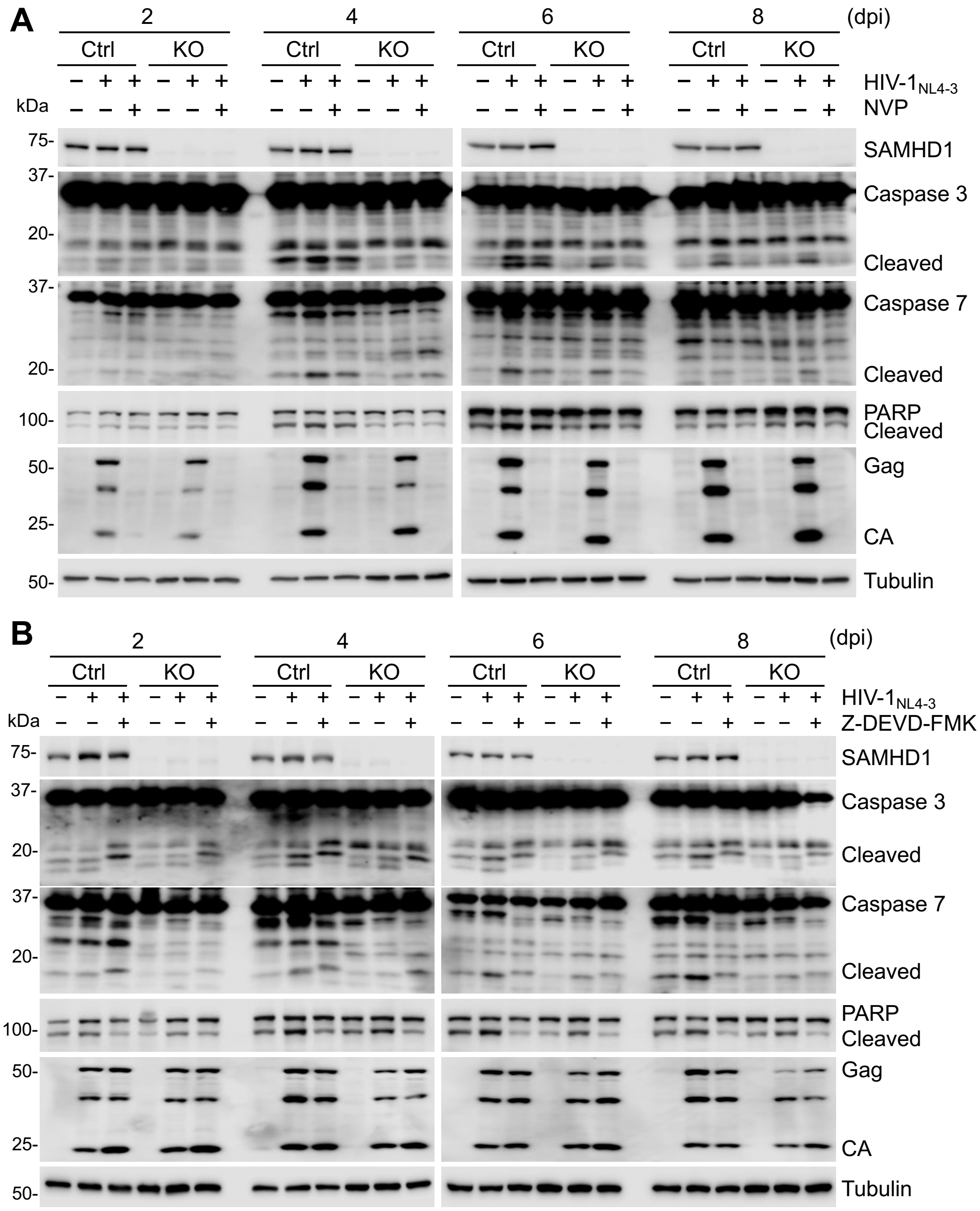


**Figure S3. NVP or Z-DEVD-FMK treatment inhibits SAMHD1-enhanced apoptosis induced by infection with replication-competent HIV-$1_{NL4-3}$**

**(A)** THP-1 Ctrl and SAMHD1 KO cells lines were infected with HIV-$1_{NL4-3}$ (MOI=2) or mock infected, and harvested at 2, 4, 6 and 8 dpi for Western blot. NVP was used to block HIV-1 replication.
**(B)** THP-1 Ctrl and SAMHD1 KO cells lines were infected with HIV-$1_{NL4-3}$ (MOI=2) or mock infected, and harvested at 2, 4, 6 and 8 dpi for Western blot. Z-DEVD-FMK was used to inhibit caspase 3/7 activity.
**(A and B)** Detection of SAMHD1, caspase 3, caspase 7, PARP, HIV-1 Gag and CA, and tubulin by Western blot.

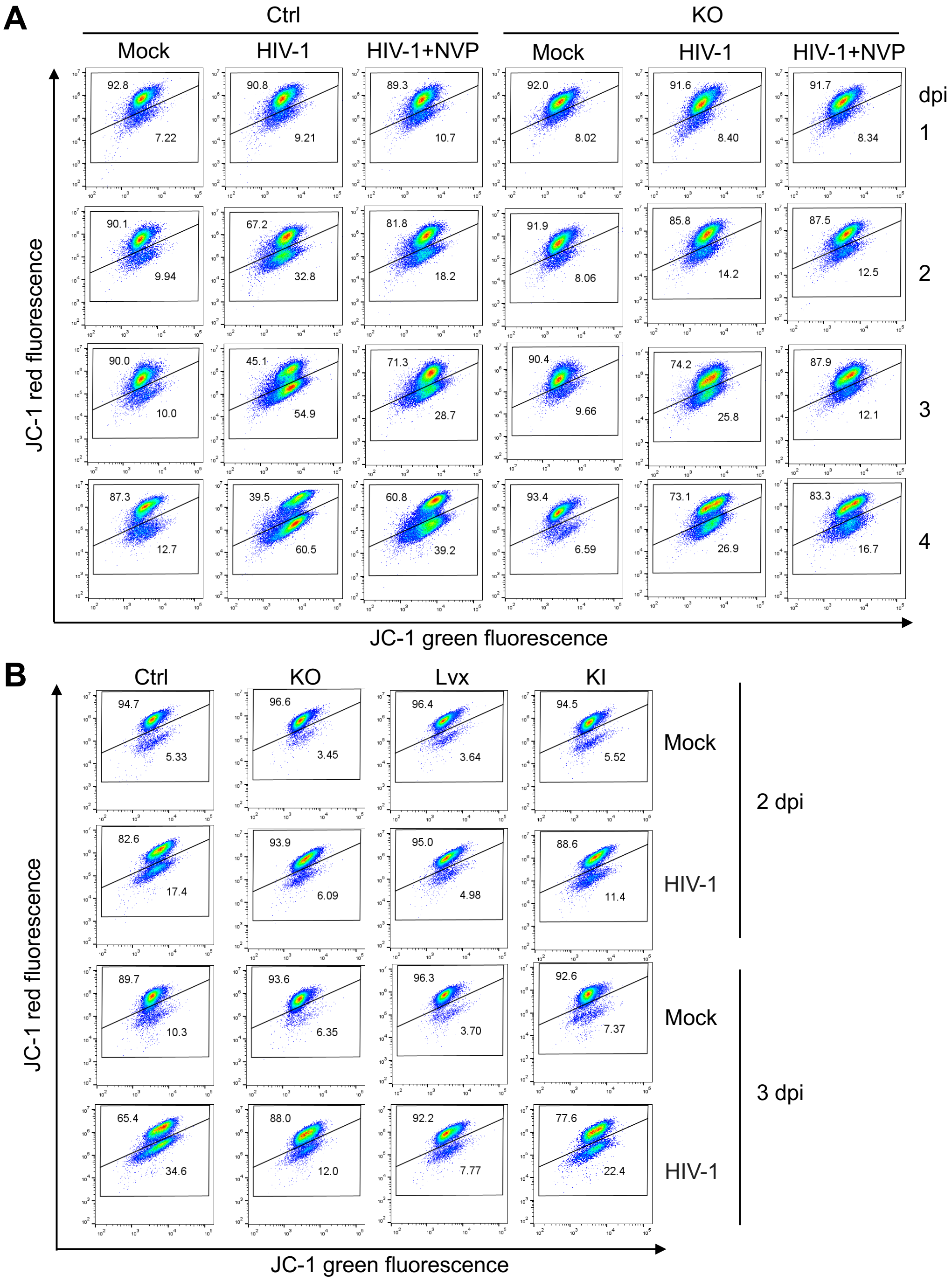


**Figure S4. SAMHD1 enhances apoptosis induced by HIV-1 infection through the mitochondrial pathway**

**(A)** THP-1 Ctrl and SAMHD1 KO cell lines were infected with HIV-1-Luc/VSV-G (MOI=2) for the indicated times or mock infected. NVP was used to block HIV-1 infection. Cells were harvested at 1, 2, 3, or 4 dpi for flow cytometry.
**(B)** THP-1 Ctrl, SAMHD1 KO, Lvx and SAMHD1 KI cell lines were infected with HIV-1-Luc/VSV-G (MOI=2) or mock infected. Cells were harvested at 2 or 3 dpi for flow cytometry.
**(A and B)** Cells were stained with JC-1 and measured by flow cytometry.

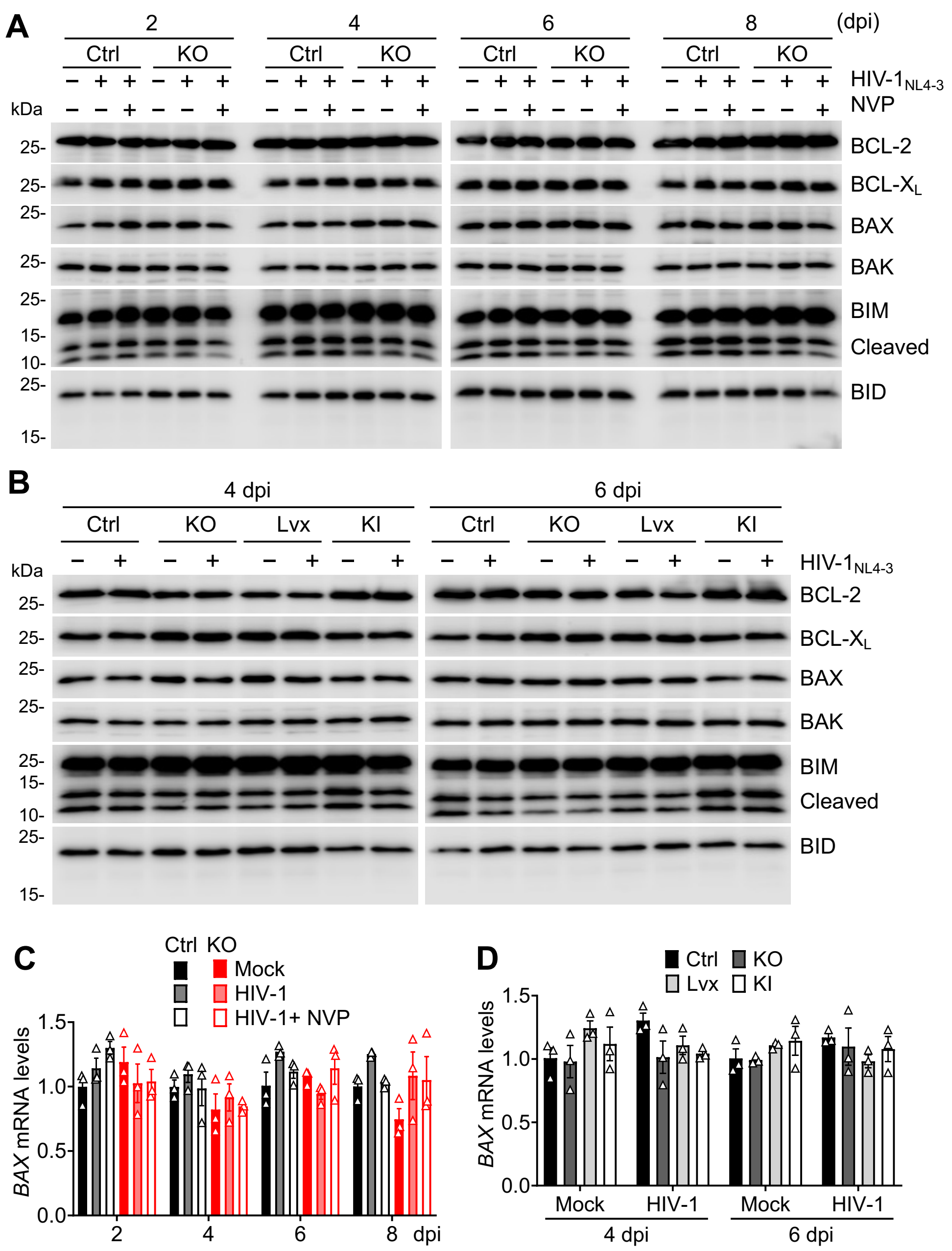


**Figure S5. SAMHD1 does not affect the expression of BCL-2, BCL-X$_L$, BAX, BAK, BIM, and BID**

**(A and C)** THP-1 Ctrl and SAMHD1 KO cell lines were infected with HIV-1$_{NL4-3}$ (MOI=2) or mock infected, and then harvested at 2, 4, 6, and 8 dpi for further analyses. NVP was used to block HIV-1 infection.
**(B and D)** THP-1 Ctrl, SAMHD1 KO, Lvx vector control, and SAMHD1 KI cell lines were infected with HIV-1$_{NL4-3}$ (MOI=2) or mock-treated, and then harvested at 4 or 6 dpi for further analyses.
**(A and B)** Detection of BCL-2, BCL-X$_L$, BAX, BAK, BIM, and BID by Western blot.
**(C and D)** The mRNA levels of *BAX* were measured by qRT-PCR and were expressed relative to mock-infected THP-1 Ctrl cells that was set to 1. *GAPDH* was used to normalize qRT-PCR results.

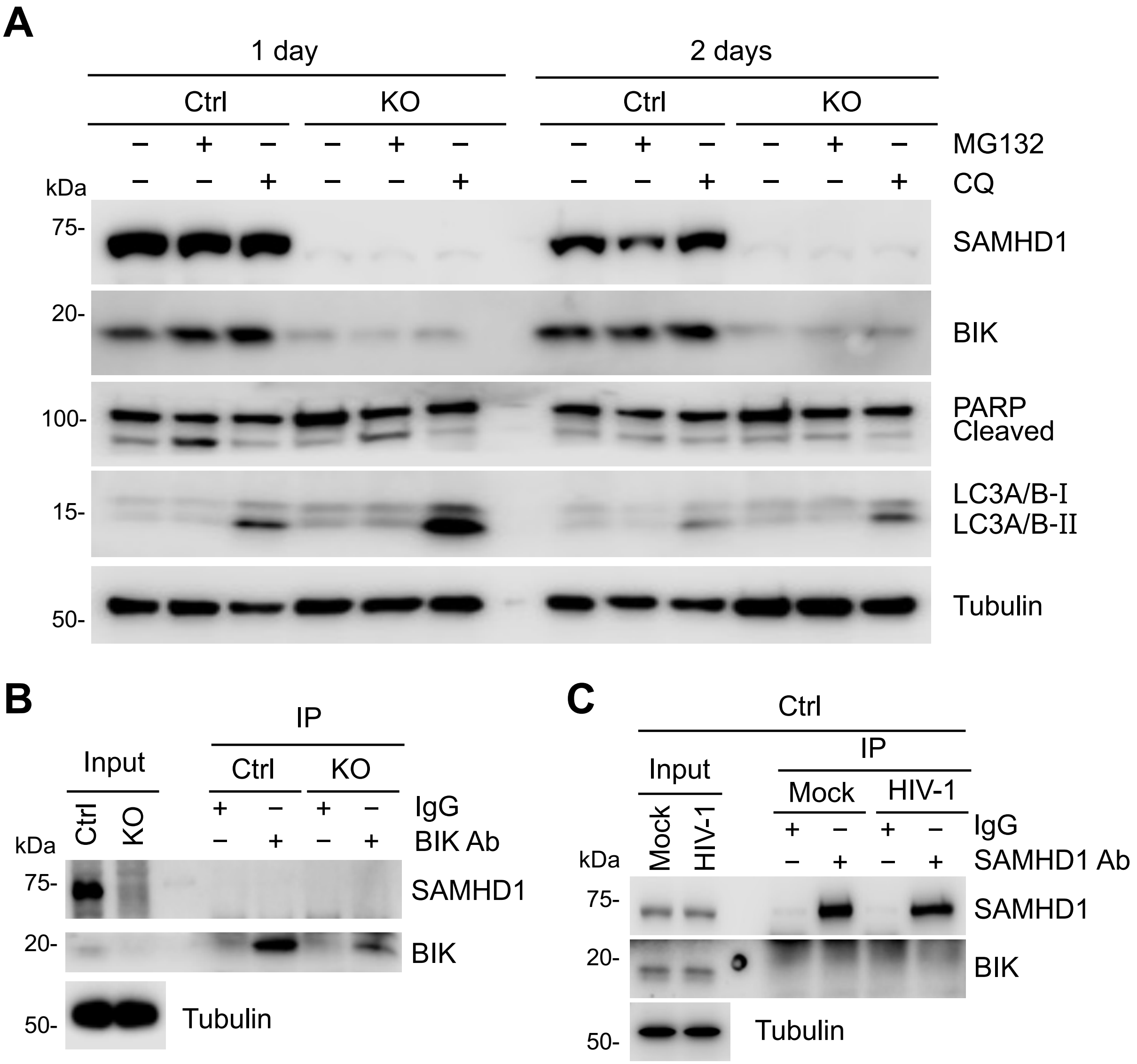


**Figure S6. SAMHD1 does not affect BIK expression via the proteasome or lysosome and does not interact with BIK**

**(A)** THP-1 Ctrl and SAMHD1 KO cell lines were treated with MG132 (1 μM) or chloroquine (CQ, 100 μM) for 1 day or 2 days, or mock treated. Detection of SAMHD1, BIK, PARP, LC3A/B, and tubulin in whole cell lysates by Western blot. Tubulin was used as a loading control.
**(B)** THP-1 Ctrl and SAMHD1 KO cell lines were lysed for immunoprecipitation (IP). BIK antibody (2 μg per $1x10^7$ cells) was used for IP. The same amounts of nonspecific rabbit IgG were used as a negative control.
**(C)** THP-1 Ctrl cells were infected with HIV-$1_{NL4-3}$ (MOI=2) or mock infected and then lysed at 6 dpi for IP. SAMHD1 antibody (2 μg per $1x10^7$ cells) was used for IP. The same amounts of nonspecific mouse IgG were used as a negative control. Input and IP products were detected by Western blot.

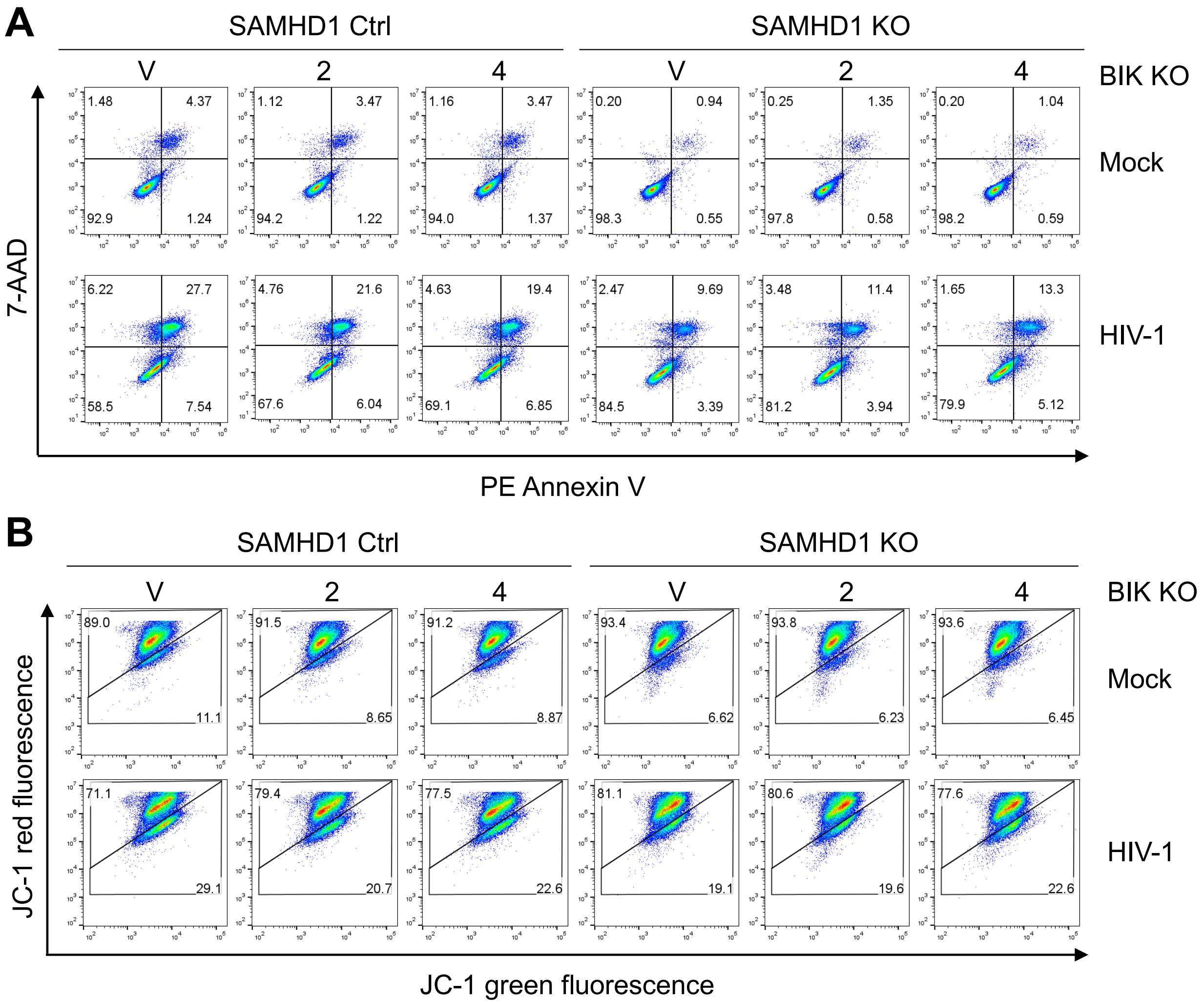


**Figure S7. Endogenous BIK contributes to SAMHD1-enhanced apoptosis induced by single-cycle HIV-1 infection**

**(A and B)** THP-1 Ctrl and SAMHD1 KO cell lines expressing empty guide RNA vector (V), BIK-specific guide RNA 2 (BIK KO-2), BIK-specific guide RNA 4 (BIK KO-4) were infected with HIV-1-Luc/VSV-G (MOI=2) or mock infected. Cells were harvested at 3 dpi for staining and flow cytometry.
**(A)** Cells were stained with Annexin V and 7-AAD and measured by flow cytometry.
**(B)** Cells were stained with JC-1 and measured by flow cytometry.
